# Automated plankton monitoring suggests a key role of microzooplankton and temperature for predicting dynamics of phytoplankton size classes

**DOI:** 10.1101/2024.02.23.581723

**Authors:** Stefanie Merkli, Ewa Merz, Marta Reyes, Pinelopi Ntetsika, Stuart R. Dennis, Peter D.F. Isles, Sreenath Kyathanahally, Marco Baity-Jesi, Sze-Wing To, Agostino Merico, Francesco Pomati

## Abstract

The interplay between abiotic (resource supply, temperature) and biotic (grazing) factors determines growth and loss processes in phytoplankton through resource competition and trophic interactions, which are mediated by morphological traits like size. Here, we study the relative importance of grazers, water physics and chemistry on the daily net accumulation rates (AR) of individual phytoplankton from natural communities, grouped into six size classes from circa 10 to 500 μm. Using a Random Forest modelling approach and four years of daily data from a lake, we find that water temperature is generally a pivotal control of all phytoplankton AR. At the same time, nutrients and light are important for the smallest and the largest classes. Mesozooplankton abundance is a key predictor of the AR for small phytoplankton, with microzooplankton being important for the middle-size range. In our data, large and small phytoplankton have different (seasonal) blooming patterns: small forms are favoured by low temperature and grazing, and high phosphorus levels. Larger forms show positive ARs at high temperatures and low phosphorus (being relatively insensitive to zooplankton grazing). These results help us understand the opportunities and limitations of using size to explain and model phytoplankton responses to biotic and abiotic environmental change.

## Introduction

Understanding the relative importance of biotic and abiotic controls of biodiversity change, in terms of taxa and traits, is critical in times of global and local anthropogenic impact on ecosystem processes. Phytoplankton in particular span enormous phylogenetic and functional diversity and regulate essential global-scale processes and elemental cycles, such as those of carbon and nutrients [1]. The relative importance of nutrient supply and trophic controls driving phytoplankton community dynamics can vary depending on the environmental conditions [2, 3], and affect large-scale ecosystem properties and processes [4, 5]. Recent studies suggest the importance of quantifying both division and loss processes or their net outcomes when studying the environmental drivers of phytoplankton community dynamics (Behrenfeld and Boss 2018; Arteaga et al. 2020). While temperature and nutrient supply mostly regulate growth processes [6], trophic controls (direct negative effects on net growth and indirect facilitation [7]) play a role in growth and loss, explaining grazing defence traits and variation in community structure [8–11]. The net rate of accumulation (AR) represents the balance between phytoplankton division rates and the sum of all loss rates (e.g., grazing, sinking, parasites), and it is essential for understanding controls of phytoplankton community diversity and for algal bloom prediction [6, 12, 13]. The AR of phytoplankton is, however, dependent on their ecological context: the outcomes of abiotic and biotic interactions are fluctuations in the daily AR of taxa [12, 14], which depend on ecological and physiological traits, and local environmental conditions [6, 12].

Morphological characteristics of phytoplankton, like the size of cells and colonies, are central traits linking the principal axes of resource competition and trophic interactions [6, 15, 16]. Realised size of organisms scales with species physiology and ecology, therefore with population growth and loss rates (i.e. AR), determining emergent properties of the whole food web [5, 6, 15]. Larger cell or colony size increases resistance against grazers, reduces the competitive ability for nutrient uptake, and increases sinking losses in stable water columns. Small cells generally possess lower sinking rates and higher maximum growth rates and should be favoured under low nutrient conditions. Under higher temperatures, small cells can maintain higher metabolic and growth rates than larger phytoplankton because they are better nutrient competitors due to their higher surface-to-volume ratio [16]. There are however uncertainties regarding these general and theoretical expectations, emerging from complex ecological interactions and variable environmental conditions. For example, spatial data in phytoplankton size-abundance relationships in lakes across the USA showed that communities do not show theoretically expected patterns based on local environmental conditions [17], while long-term responses of phytoplankton size-abundance relationships in Swiss lakes suggest that small forms may be favoured under high resource levels and low temperatures, whereas large (often mixotrophic) taxa prevail under low resource conditions and higher temperatures [17].

Resolving the above incongruencies would be important to understand the effects of climate change and eutrophication on plankton communities and aquatic ecosystem processes. Understanding the relative importance of biotic and abiotic controls on macroecological patterns such as size distributions in natural communities remains however a challenge. First of all, it is difficult and time-consuming to manually estimate the realised size of individual plankton organisms, which is a plastic trait (variable over space and time), as opposed to the nominal size of species (which is unrealistically fixed). Additionally, measuring ecologically relevant ARs would require monitoring of phytoplankton abundances at biologically relevant timescales, such as days (i.e. the generation time of the focal organisms), which is intractable with traditional monitoring methods. Lastly, to study the relative importance of abiotic and biotic factors influencing phytoplankton ARs, high-frequency monitoring of physicochemical water variables and zooplankton taxa would also be necessary.

The purpose of this study was to investigate the temporal prevalence, relative strength and effect of (i) available resources (phosphorus, nitrogen, light), (ii) water physics (temperature, turbulence, stability), and (iii) herbivore grazers (daphnids, copepods, ciliates, rotifers), in controlling the daily AR of phytoplankton size-classes spanning the range between circa 10 to 500 μm. We use a unique dataset of four years of daily automated *in-situ* imaging data of phytoplankton, zooplankton, and water physicochemical parameters from a freshwater lake. We study ARs with a non-parametric machine-learning approach, i.e. Random Forest (RF) modelling, which allows us to overcome several limitations of statistical methods when dealing with large, high-dimensional datasets [19].

Specifically, we test the following hypotheses:

- Smaller phytoplankton forms are favoured (i.e. have higher AR) under low resource conditions, and/or higher temperature and water column stability.
- Larger phytoplankton require higher resource levels than smaller forms, and/or have a competitive disadvantage under higher temperatures (low AR), and are more susceptible to sinking losses under a stable water column.
- Smaller phytoplankton forms are more strongly controlled by herbivores (low AR) whereas large phytoplankton are more resistant to (size selective) grazers (high AR).

## Materials and Methods

### Plankton data collection/processing

We used high-resolution monitoring data to study the plankton community in a lake in Switzerland (Lake Greifensee) between May 2019 and June 2023. Greifensee is a lowland morenic lake in the Swiss pre-alps, naturally meso-eutrophic (max. depth 32 m, average depth 18 m), with a documented history of anthropogenic eutrophication, which was reverted by controlling phosphorus releases in the 1980s-90s [4]. Greifensee completely mixes once per year in winter and, at circulation, has a load of 0.04 mg/L of total P, which makes it Eutrophic according to the OECD guidelines, and a load of 1.25 total N (https://hydroproweb.zh.ch/Karten/JB%20Chemie%20Seen/Dokumente/02_Be.pdf).

Plankton was monitored with a dual-magnification dark-field imaging microscope that captures all particles in the size range between ∼10 μm and ∼ 1 cm at 3 m depth using two magnification objectives: one for phytoplankton and small zooplankton (5.0x) and one for larger zooplankton (0.5x) [20, 21] (**Fig. 1A**). The choice of depth is motivated by the average structure of the lake water column during the growing season: the thermocline is generally around 8 m (https://www.datalakes-eawag.ch/datadetail/515) and the epilimnion is generally well mixed by daily thermal winds. The 3 m depth is generally a representative sample of the phytoplankton community of the photic zone. Images were collected at a rate of 1/s for 10 minutes every hour and regions of interest (ROIs) were automatically identified by the instrument using a Canny edge detector [21]. A Python image processing script performed colour conversion, edge-detection, and morphological feature extraction (e.g. object dimensions) of detected raw ROIs (https://github.com/tooploox/SPCConvert). Objects were manually annotated and zooplankton was finally classified into taxa using an ensemble of trained deep-learning classifiers (convolutional networks) [22, 23]. We removed small zooplankton from the 5.0x data and estimated body size as the area of the binary image mask for each phytoplankton object [20]. Such body size estimates scale with traditional microscopy biovolume measurements [21].

**Fig 1:**
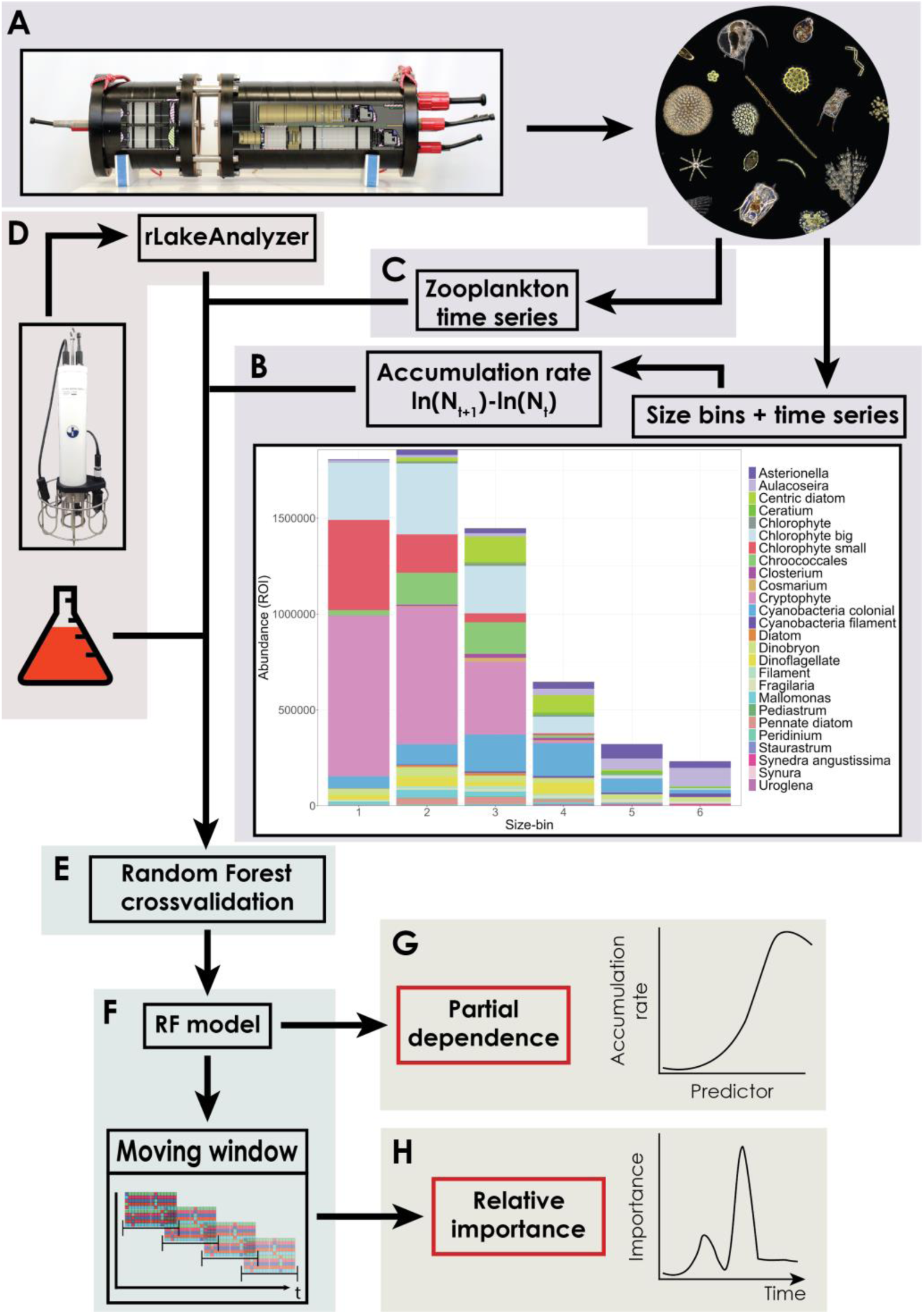
General workflow of the study, summarising i) the type of input data, both biotic (A-B-C) and abiotic (D), ii) the data analysis approach (E-F) and iv) the modelling outcomes (G-H). Specifically: **A**: Images of phytoplankton and zooplankton from the underwater microscope: size and taxonomic classification are obtained by image processing and deep learning, respectively. **B**: Phytoplankton is assigned to 6 size-bins based on the area (as a measure of body size) of each imaged object (**Tab. S2**); daily accumulation rates (AR) are calculated based on Log-abundances. **C**: Mesozooplankton (daphnids, cyclopoid and calanoid copepods) and microzooplankton (nauplii, rotifers and ciliates) daily time series are also obtained from (**A**) (**Table S2)**. **D**: Water physicochemical variables are extracted from high-frequency CTD profiling data, whereas daily nutrient chemistry is predicted from weekly laboratory measurements (**SI Methods**). **E**: Random Forest (RF) models are validated, and used to study the relative importance and temporal prevalence of AR predictors with a rolling-window approach (**F-H)**. **G**: Partial effects of important AR predictors are extracted from the trained RF models.

To divide the phytoplankton community into size bins, we temporarily excluded the first and last percentile of the size distribution before binning to give the tails of the distribution less weight. We then binned the remaining objects into 6 bins of equal length on a Log_10_ scale (bin function from the OneR package V2.2) and returned the first percentile to the first bin and the last percentile to the last bin (**Fig. 1B, Tab. S1**)[24]. Note that individual phytoplankton cells or colonies are assigned to different bins irrespective of their taxonomic identity, and solely based on each object size (**Fig. S1**). We aggregated (summed) hourly abundances per day per size-bin: abundances are in ROI/sec (region of interest per second), which correlates with plankton concentrations [21]. We used LOESS regression to smooth the time series with a degree of smoothing of α = 0.025. We chose α to exclude only random (temporal non-autocorrelated) noise. We calculated daily accumulation rates (AR), for each bin, as:

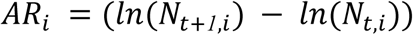

Where i = size-bin, N_t_= abundance at time t and N_t+1_ = abundance at time t+1.

As a measure of trophic controls on phytoplankton AR, we estimated the daily abundance (summed of hourly ROI/sec) of daphnids, cyclopoid copepods, calanoid copepods, nauplii and rotifers (from 0.5x), and ciliates (from 5.0x) (**Fig. 1C**, taxonomic composition of classes in **Tab. S2**). We scaled zooplankton abundances to z-scores for modelling.

### Abiotic data collection/processing

Lake physical and chemical variables were automatically measured four to eight times per day using a CTD probe and profiler (www.idronaut.it). In addition, a meteorological station on the roof of the monitoring station collected data continuously (Vaisala Oyj WXT520 & OTT HydroMet WS700-UMB). Once per week, in the same location, water samples from 3 m depth, near the underwater microscope, were analysed for nutrient chemistry (P-PO_4_, N-NO_3_, NO_2_ and NH_4_) (**Fig. 1D**).

As abiotic predictors in the AR models and measures of water column physical structure, we calculated thermocline depth, mixed layer depth, epilimnetic temperature using the R package rLakeAnalyzer V1.11.4.1 [25] and CTD data from the monitoring station [26]. For the estimation of all variables, we used the temperature profiles generated by the CTD probe. For the estimation of the epilimnetic temperature, we used lake topology (depth and the lake area at each depth) in addition to the temperature profiles. If no thermocline was detected using a minimum density gradient of 0.1 m, we used site depth as thermocline depth. We calculated daily averaged values for all variables described above.

As a proxy for light availability available to the phytoplankton community, we extracted the depth where photosynthetically active radiation (PAR, wavelengths 400-700 nm) reached 5 μEm^−2^s^−1^ (the instrument’s detection limit/sensitivity), which we call light penetration depth and depends on incoming irradiance and water turbidity (**Fig. S2**). For this, we excluded all PAR measurements from the surface to the depth of maximum PAR, where the PAR sensor was shaded by the automated monitoring platform above. We then reported the depth at which a PAR value of 5 5 μEm^−2^s^−1^ was found (**Fig. S2**).

Measurements of the inorganic nutrients nitrate (NO_3_), ammonium (NH_4_) and phosphate (P-PO_4_) were performed weekly. To obtain nutrient levels at a daily frequency, we predicted daily values using a Random Forests (RF) model trained and validated for each nutrient (**SI Methods**). Nutrient levels fluctuate on a daily scale mostly due to variations in water physical parameters and meteorological conditions [27]. Interpolation of nutrient values based on autocorrelation of the time series would not account for such a daily variation. We therefore trained RF models on weekly nutrient data (n = 201) based on CTD and Meteo information from the same date, tested and validated these nutrient models on unseen nutrient values (10-times cross-validation where 80% of the data were used for training and 20% for testing), and then used RF models to predict (as opposed to interpolate) daily nutrient concentrations based on daily data of water physics and meteorological conditions (**Fig. 1D**). For more information nutrient models and their performance see **SI Methods**, **Table S4** and **Fig. S12**.

### Random Forests for Modeling Complex Relationships

To extract robust patterns from our complex and large datasets we used RF [28, 29], which allowed us to overcome the most important constraints of traditional statistical approaches: a priori specification of (i) functional forms for partial effects of explanatory variables, (ii) interactions and non-linearities, and (iii) error distributions (**Fig. 1E**). By bootstrapping both data and variables, RF are relatively robust to collinearity. To reduce the tendency of these models to overfit the data, we smoothed the response variables (AR, see above).

### Size bin Modelling

We modelled daily ARs of size bins (change in Log abundance between time t and t+1) as a function of the abundance of each size bin (the density-dependence of each bin’s growth) and various abiotic and biotic explanatory variables, all at time t (i.e. lagged by 1 day), to capture contemporary (i.e. direct) effects of environmental conditions on phytoplankton dynamics. This allowed us to model the process of net (and proportional - given the Log scale) accumulation of phytoplankton in each size bin by accounting for the density dependence of population growth, and the focal biotic and abiotic mechanisms. By focusing on these factors measured at the same time point, we targeted the mechanisms directly affecting daily phytoplankton dynamics, such as organismal division and loss. This approach is particularly suitable considering the short time scale (daily) of these processes. We acknowledge that lagged effects of environmental variables can also influence phytoplankton abundance, but these indirect and time-delayed interactions were not the focus of this study. Our RF approach, based on a rolling-window analysis of size-bin time series (**Fig. 1F**), describes how the predictors of AR vary in their relative strength (i) over time (within size-bins) and (ii) between size-bins (**Fig. S6**). While the former can be expected in nonlinear dynamic and chaotic systems [31–33] and will not be discussed in detail, aggregating the relative importance of predictors over time allows us to study the variation between different size-bins (**Fig. 1H**). Examining the partial effect of environmental predictors when the RF model is trained on all data (**Fig. 1G**), allows us instead to extract patterns of how size-bins’ ARs relate to the different abiotic and biotic environmental factors.

### Random Forest training & validation

We used RF models to estimate the relative importance of each explanatory variable in predicting phytoplankton AR per size-bin and their effects as partial dependence plots [19]. We imputed missing values (2.7%) among predictors with the function rfImpute and then trained a RF model for each size-bin, with the randomForest package V4.6.14[30]. To rule out overfitting, we performed a random split ten-fold cross-validation, which was repeated ten times. We extracted the R^2^ from a linear model fit between the observed and the predicted accumulation rate from an RF model based on 90% of the data. By comparing the performance in the training data with the validation data, we assessed how well the models generalise to unseen data. Once we assessed that our RF models performed well and did not show overfitting, we fit a model on the entire time series (**Fig. 1E, Fig. S3**). These bin-specific RF models yielded pseudo-R^2^ values of circa 60% (**Tab. S1**) [31]. We refer to the explanatory variables of our RF models as predictors, given the underlying method by which their importance is evaluated (loss of predictive skill of the model when the variable is omitted), without implying that patterns indicate causal links between explanatory and response variables.

### Rolling-Window Analysis for Temporal Importance

To study both the temporal prevalence and the relative strength of AR predictors, we used a rolling-window approach. Within a temporal window of 28 days, shifting forward by one week at each round, we predicted the AR using the trained RF (see above) from which we calculated the increase in mean-squared-error (IncMSE) after randomly resampling (reshuffling with replacement) the vector of values of each predictor in sequence (**Fig. 1F**). Specifically: within each window, we predicted the AR with our RF model based on the entire time series. We then fit a linear model between the observed and the predicted AR. We calculated the mean squared error (MSE) as:

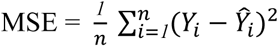

where n = number of observations, (*Y*_*i*_ − *Ŷ*_*i*_) = residual errors of each data point in a linear model of observed vs predicted AR, *Y*_*i*_ = ith true data point and *Ŷ*_*i*_ = ith estimated data point.

In this process, for each explanatory variable, we disrupted the relationship of the given variable with all other variables by shuffling the values of the given variable inside the rolling window. We again predicted the AR and calculated the MSE. We then calculated the increase in MSE (IncMSE).

Given the complexity of models (based on 13 variables) and the frequency (information content) of data, the IncMSE emerging from shuffling variables’ vectors (as opposed to omitting the variable) can be relatively small. Omitting variables would however change the model structure, making comparisons between models lacking different variables unfair. IncMSE in our approach is only used for comparisons between variables, with predictors with high IncMSE representing relatively more important variables for the model. This relative importance of predictors (IncMSE) was then reported and averaged over time, for each size bin.

### Partial Effects on AR

To study the partial effects of important predictors on AR, we trained an RF model with the whole dataset (to gain more confidence on inferred patterns) for each size-bin, and extracted as partial dependence plots of AR to the focal variables with the pdp package, V0.7.0[32] (**Fig. 1G**, **SI Methods**). All data analyses were performed with R, V4.1.1 (R Core Team 2021), and figures were produced using the package ggplot2, V3.3.5 (Wickham 2016).

## Results

The dynamics in the abundance of phytoplankton size-bins follow roughly the seasonal blooming succession of a temperate eutrophic lake, with peaks in spring, a trough in May (the clear water phase), and peaks in summer and early autumn (**Fig. 2A**). Each year, however, shows slightly different patterns in the intensity and number of phytoplankton peaks[33]. For example, the smallest size-bins are on average more abundant than the larger ones (**Fig. 2A**), as expected, except for winter 2022/2023 where the largest size-bin was dominant due to a very uncommon and persistent bloom of the diatom *Aulacoseira sp.* (**Fig. S4**).

**Fig. 2.**
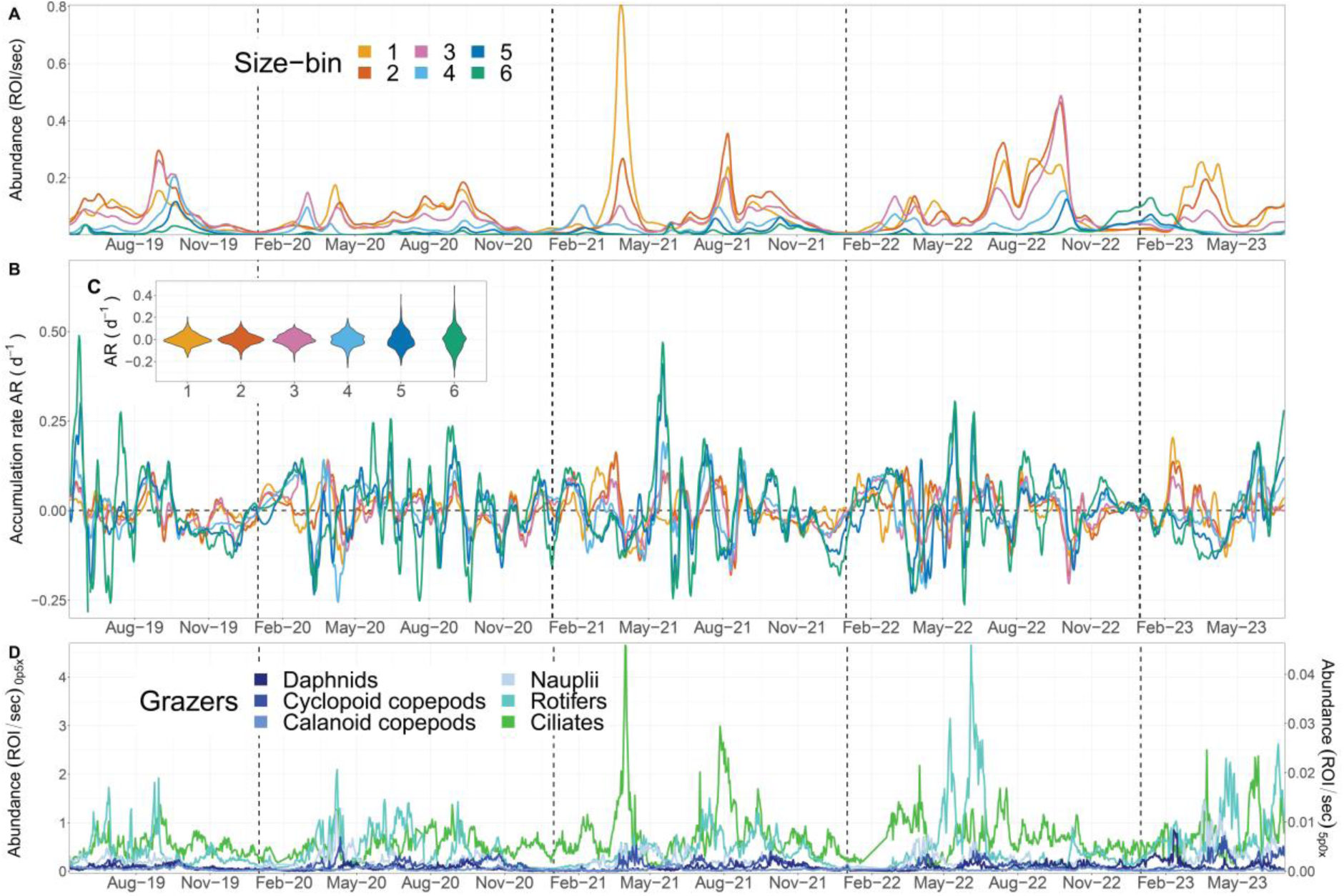
Temporal dynamics of plankton in Greifensee (2019-2022). Vertical dashed lines mark January 1^st^ of each year. **A)** Daily abundance data of phytoplankton size-bins (smoothed, see Methods). **B)** Daily accumulation rate (AR) of size-bins. **C)** Distribution of AR in each size-bin time series. **D)** Daily zooplankton abundance data (ciliates are on the right-side Y-axis because they were monitored through the 5.0x objective).

Coupled with seasonal changes in abundance, the daily process of net accumulation (AR) of each phytoplankton size-bin fluctuates over time highlighting the start and the end of the blooming periods (**Fig. 2B**). ARs vary substantially within and between size-bins, with larger phytoplankton showing a wider range of positive and negative ARs (**Fig. 2C**). Large phytoplankton do not show more rapid bloom dynamics compared to small ones (**Fig. 2A**). Among the herbivore grazers, whose trends follow the phytoplankton seasonal succession, rotifers and ciliates are the most abundant and dynamic groups (**Fig. 2D**)[34]. Abiotic environmental conditions in Greifensee follow the seasonal pattern of a temperate and monomictic eutrophic lake and are reported in **SI Fig. S5**.

The results of modelling the daily AR of each size-bin (**Fig. 2B**) as a function of their abundance and the focal abiotic and biotic explanatory variables (**Fig. 2C**, and **SI Fig. S5**) highlight important general patterns and notable size-specific differences. Aggregating the relative importance of variables over time, we find that water temperature is among the most important predictors across all phytoplankton (**Fig. 3**). The relationships between AR and temperature however, which emerge from the RF predictions of its partial effect, differ among size-bins (**Fig. 4**). Size-bins also vary in sensitivity to levels of nutrients and different grazers (**Fig. 3-4**). ARs of all size-bins show to have a similar response pattern to water mixing (mixed layer depth and thermocline depth - **Fig. S7-S8A**), which may indicate that the optimal depth of the photic zone (to balance light penetration and nutrient supply from the hypolimnion) is between 4-8 m for all phytoplankton in the lake [38]. We see that ARs saturate with increasing light intensity (depth at 5 PAR), as expected [19], and that larger phytoplankton have a higher AR at high light levels compared to smaller forms (**Fig. S7**). Size-bins display slight differences in the optimum levels of ammonia. The correlation between AR and phosphate variation is instead antithetical between small and large size-bins (**Fig. 4**): large phytoplankton show high AR at low phosphorus concentrations, while small forms peak in AR (and saturate) at high levels.

**Fig. 3.**
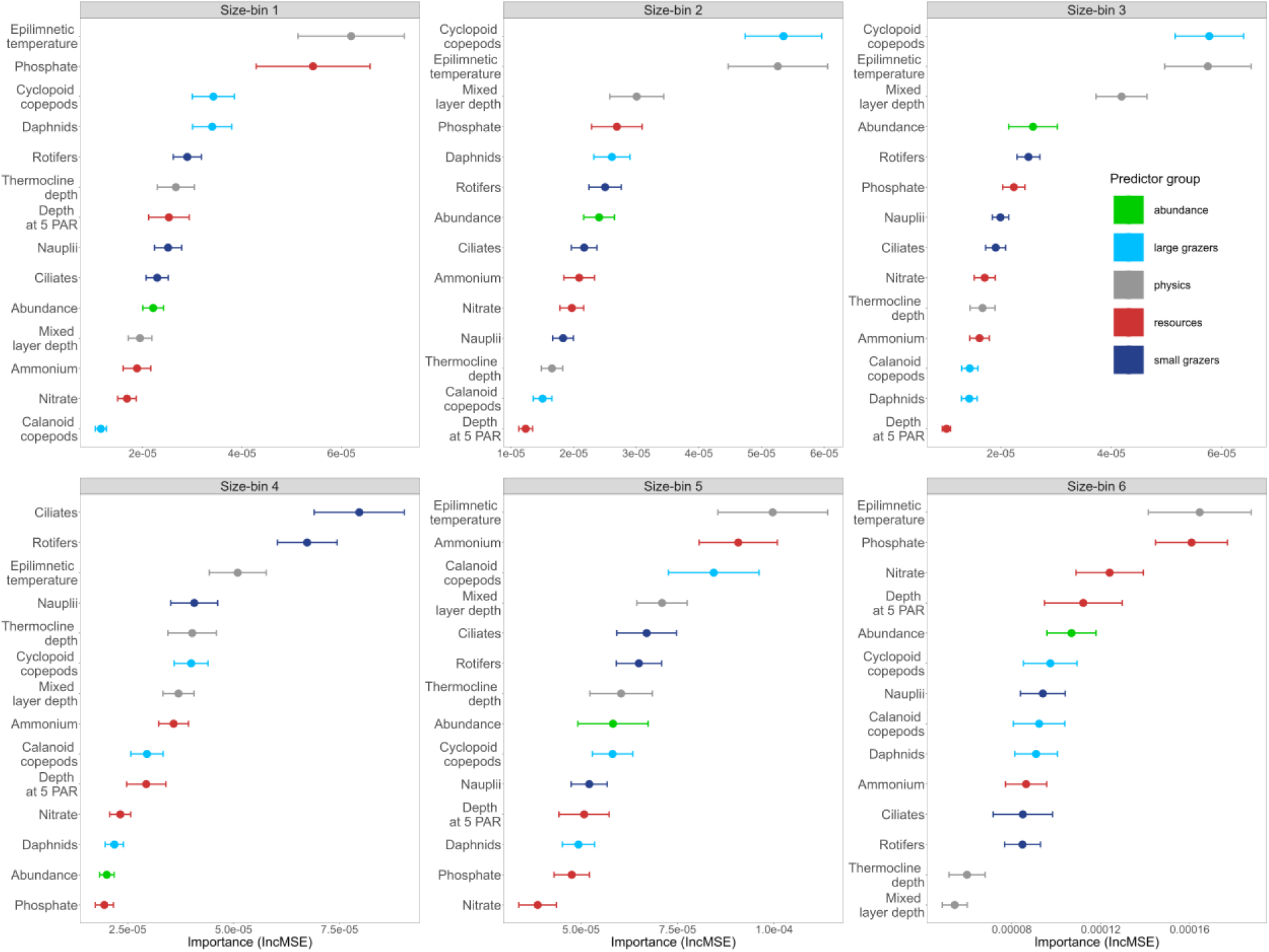
Overall mean importance (IncMSE ± standard error) of each environmental variable over the entire study period for each size-bin. Colours indicate types of predictors: grey = water physics, red = resources, light blue = mesozooplankton, dark blue = microzooplankton, green = size-bin abundance.

**Fig. 4.**
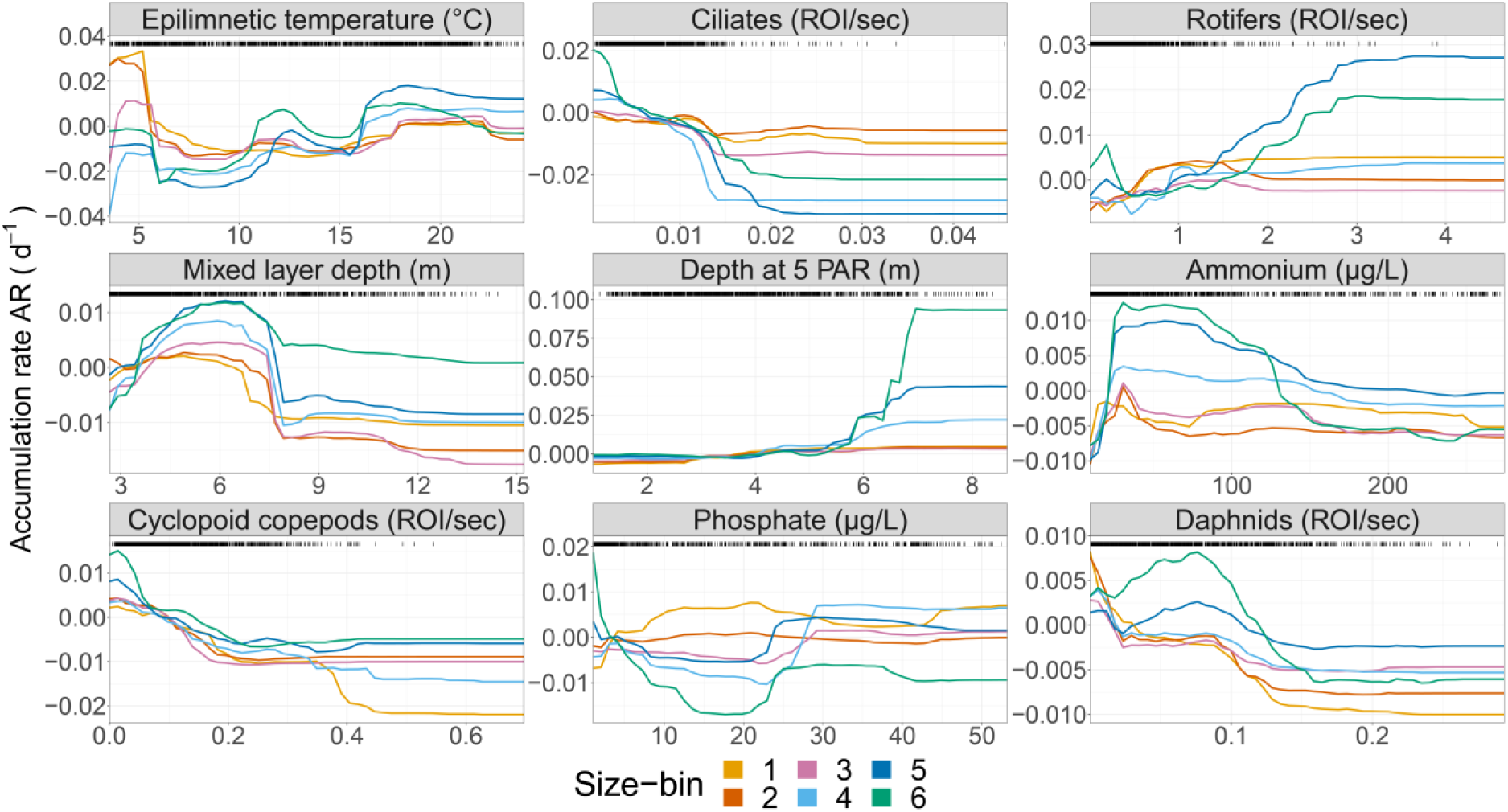
Partial effects of selected environmental variables from RF model predictions (for all of them, see **Fig. S7)**: these dependencies emerge from predicting the response (AR) over varying levels of the chosen predictor (X-axis, in the plot title) while holding the other variables in the RF model at their medians over the entire dataset. Each line represents one size-bin (colours as in Fig. 1). The distribution of observed data points is shown as black bars on top of each graph: high data-dense areas hold more confidence in the prediction.

We find that the AR of smaller size-bins is related more strongly with water temperature, phosphate, and herbivore grazing by mesozooplankton (**Fig. 3**). Particularly, the AR of size-bins 1-2 shows a peak of positive AR for low water temperature and cyclopoid copepod abundance, and phosphate in the range between 15-30 μg/L. Smaller phytoplankton also seems to have higher ARs at intermediate levels of light penetration (depth at 5 PAR) (**Fig. 4**). This is suggestive of a springtime blooming environment (**Fig. S5**), which is characterised by communities of small diatoms, green algae and cryptophytes (bins 1-2, **Fig. 1**) [34]. Conversely, larger size-bins (i.e. 5-6, characterised by large colony-forming taxa, including cyanobacteria) were shown to be less controlled by grazers, compared to the other groups, and to have optima at high temperature and light penetration, and low phosphate levels, which are typical of summer conditions (**Fig. S5**).

Increasing density of cyclopoid copepods shows negative effects on the AR of size-bins 1-4, suggesting trophic control (herbivory) on small and medium phytoplankton (**Fig. 4**). Similarly, daphnids also show generalised negative effects on ARs, and both daphnids and cyclopoids affect ARs in a nonlinear way that suggests some form of indirect facilitation on small and large phytoplankton forms, respectively, at high density (**Fig. 4**). Rotifers, which are important predictors for small to medium-sized phytoplankton, have a similarly nonlinear and slightly positive effect on ARs (**Fig. 3-4**). Ciliates are relatively more important as predictors of medium-sized phytoplankton AR (bins 4-5), with the expected negative effect of herbivore grazing (**Fig. 3-4**). We note that, in general, and despite differences in their effects, microzooplankton is a very important biotic predictor of AR across all size-bins, but particularly for intermediate size. The largest size-bins (5-6) seem to be relatively less affected in their ARs by the density of zooplankton, the most important being, for these size classes, calanoid and cyclopoid copepods (**Fig. 3-4**).

## Discussion

In this study, we leverage a unique dataset of four years of automated plankton imaging data to investigate whether size-based phytoplankton groups respond to environmental conditions as expected from theory, experiments, and observations at coarser time scales [6, 15, 16]. This study is different from previous ones in four aspects. First, while body size derived from imaging individual organisms depends on some image processing assumptions [39, 40], with high-frequency measures of millions of individual plankton organisms across the entire community (from 10-500 μm) we capture variation in this trait associated with inter and intraspecific processes (physiology, ecology and, possibly, evolution) [41]. Second, having high-frequency data, we study dynamics at short time scales by modelling daily net accumulation rates of size-bins (a growth process), not their abundance (a state variable). Third, we model AR of size-bins based on abiotic and biotic variables measured *in situ*, the latter comprising virtually all of the functional components of the plankton community. Lastly, we use machine learning to reduce modelling assumptions to the minimum and to extract robust non-linear patterns from large and noisy datasets [29]. Our RF models have an R^2^ of circa 0.6 in validation (**Fig. S3**), which suggests that we captured deterministic environmental factors that relate to size-bin dynamics: here we discuss how much these patterns resemble known size-mediated mechanisms driving phytoplankton responses to environmental conditions.

While the observed dynamics of phytoplankton size-bins follow a seasonal pattern similar to the one expected by temperate eutrophic lakes [31], we note that each year shows differences in the intensity and number of phytoplankton peaks, highlighting the intrinsic challenge of understanding plankton communities in chaotic systems where even small perturbations can propagate over time influencing dynamics [32]. Larger phytoplankton also showed a wider range of positive and negative ARs (**Fig. 2C**), probably due to the AR of bins of larger organisms, which are less abundant, being more sensitive to daily variation in numbers compared to smaller, more common phytoplankton.

In this context, we see little evidence for resources increasing in their relative importance with increasing phytoplankton size (**Fig. 3**, **Fig. S9**), as expected from the Metabolic Theory of Ecology [16, 42]. This is because, in a eutrophic lake (high nutrient levels) and in the context of the complex and non-linear biotic and abiotic interactions of a plankton community, nutrient limitation may play an important role only sporadically (**Fig. S6**). We also find weak support for larger phytoplankton being favoured by deeper mixing (mixed layer depth - **Fig. 4**): this effect, however, may be confounded by the fact that deeper mixing is generally associated with upwelling of nutrients [43]. In line with previous studies [16, 18], we find that AR of smaller size-bins (1-2, **Fig. 3**) is more strongly correlated to water temperature and herbivore grazing. Smaller phytoplankton have higher ARs at intermediate levels of light, suggesting blooms during springtime, which is characterised by communities of small diatoms, green algae and cryptophytes (bins 1-2, **Fig. 1**) [31]. We find in fact that small phytoplankton have higher AR at low temperatures, and large forms have higher AR at higher temperatures (**Fig. 4**). This has important implications for understanding and predicting algal blooms, which are generally characterised by large cells [44, 45]. Previous studies reported opposite trends [42, 46], however a similar pattern as in this study was previously reported in lakes, where summer blooms of cyanobacteria are common [18]. These differences can be attributed to the fact that species composition within size classes varies substantially over environmental gradients: specifically for this study, a large number of cold-adapted species (e.g. diatoms, cryptophytes) can be found in the small size-bins, and many warm-adapted species (e.g., colonial cyanobacteria) in the large size-bins. Cold-adapted species have lower thermal optima, which leads to a steep decrease in AR when exposed to temperatures further away from their optimum (**Fig. 4**) [6]. *Vice versa*, warm-adapted species, present in the larger size-bins, show AR that increases steadily with water temperature (**Fig. 4**, **Fig. S7**). These results suggest that the blooming of small and large phytoplankton may be triggered by different biotic and abiotic mechanisms.

One important pattern emerging from our data is that microzooplankton is an important predictor of AR for all phytoplankton regardless of size. This finding has implications for understanding the diversity of planktonic food webs and their responses to environmental change because microzooplankton are selective feeders, are highly dynamic (fast growth rate) and central to plankton interaction networks [4]. We expected microzooplankton to be selective for small phytoplankton forms [46] but, to our surprise, they appear very important for medium-sized bins (4-5, **Fig. 3**, **Fig. S9**). Even though in our study microzooplankton is composed of defined classes (nauplii, rotifers and ciliates), each one of them is a heterogeneous group of microorganisms belonging to different species and lineages, with large cryptic diversity in terms of physiology, life history, and size selectivity - all of which also depend on the composition of the available preys and environmental conditions. Microzooplankton can exhibit varying degrees of size selectivity depending on the species and the available prey - while in general, they are supposed to prefer smaller phytoplankton sizes, their actual preference can vary from a few micrometres to several tens of micrometres (as in the case of preys such as diatoms or filamentous cyanobacteria) [47–49]. Rotifers are often opportunistic omnivores, with potential trophic cascade effects (e.g., indirect facilitation) on phytoplankton by eating flagellates and ciliates [50–52]. This may explain why rotifers positively correlate with large phytoplankton in our study. Alternatively, microzooplankton may contribute to releasing or recirculating nutrients locally when they are at high density [10, 53].

Mesozooplankton seems to be a more important predictor of small phytoplankton forms [46]. Daphnids graze on phytoplankton with a size smaller than their gape, i.e., with a diameter of < 20 or 30 µm, depending on the species [54, 55]. Calanoids and cyclopoids show differences in feeding behaviour, which also depend on life stages: cyclopoids prey mostly on large particles, while calanoids consume smaller particles [56]. Some cyclopoid species are also primarily carnivorous [57]. Contrary to these expectations, cyclopoid copepods in our study are important predictors for the AR of smaller-sized phytoplankton (bins 1-3). In general, mesozooplankton are also more efficient grazers and, combined with the higher encounter rates for the smaller and more abundant phytoplankton, could cause a larger grazing impact on the abundance of smaller forms, as opposed to larger, rarer and more defended phytoplankton. This suggests that mesozooplankton may have a role in the accumulation of large and colony-forming phytoplankton, including a role in cyanobacterial bloom development [13].

The predicted effects of environmental conditions on AR in our RF models are mostly non-linear (**Fig. 4**). Some of these correlations match prior expectations from growth rate response curves obtained under laboratory conditions (e.g., responses to resources show a Monod or a Gaussian shape), others do not (e.g. temperature and P-PO_4_ with both concave and convex curves). We find that the shape of these effects varies between phytoplankton size-bins, while we were expecting them to be relatively universal [6]. One explanation for this unexpected variability is that such effects may depend on the interaction between multiple environmental factors, and show the expected patterns only in certain areas of a multi-parameter space. These interactions can be very complex and interdependent - e.g. the effects of nutrients depend on temperature (see **Fig. S10**) [4]. Similarly, also the effects of biotic factors depend nonlinearly on abiotic conditions like water temperature (**Fig. S11**), implying that unidimensional response curves are strongly context-dependent, and for understanding and predicting phytoplankton dynamics a multidimensional approach is required [58]. Alternatively, the above and unexpected patterns may be due to four important caveats of our study.

First, compositional turnover of taxa within size-bins may confound some of the detected partial effects of environmental conditions on AR (**Fig. 4**), and possibly also their relative importance (**Fig. 3**): taxa composition and relative abundance change within size-bins over time (**Fig. S12**), and different taxa have different ecology and physiology. Second, the realised size of individual phytoplankton objects, as studied here, is not the taxon cell size as intended in most of the previous theoretical and experimental work: the realised size, whether cell or colony size, is a plastic trait at the individual level (e.g. mediated by physiological and ecological interactions), as opposed to a nominal feature - not measured for each individual object, but assumed based on species average dimensions and shape - as in many previous investigations [5, 15, 17, 18]. This could have influenced some of the unexpected results regarding the effects of grazers (which may target the cell inside the colony irrespective of colony size) and of abiotic factors (e.g., temperature - large taxa are colony-forming summer species). We note that our study uses objects’ area as a measure of body size, instead of volume as in many previous studies: this difference should not however have an impact on the construction of the size-bins used here, and on the temporal dynamics of their abundances (see **SI Results**). Third, size, as acquired from imaging, shows high variation within and between taxa also due to organisms being photographed from different angles, and sometimes only partially imaged [40]. This variation is not related to responses to environmental conditions. Lastly, the dynamics of size-bins could have been mediated by cell shape. Environmental factors can influence the selection for body size and shape differently [59]: shape variation represents a competitive strategy for medium and large phytoplankton forms to acquire an advantageous surface area-to-volume ratio [60], or to reduce sinking velocity [61]. Future studies should consider including shape in trait-based modelling of size classes.

In conclusion, we find only partial support for response patterns to environmental factors in size-based phytoplankton classes, as expected from the literature. This can be due to the difference in our study compared to previous work, as explained at the beginning of the Discussion, or to the caveats mentioned above. More broadly, however, our study shows the power but also highlights some limitations of size-based approaches to study plankton dynamics. First, studying phytoplankton communities may require complementing size information, measured at the individual level as done in this study, with either shape and/or taxonomic identity, to increase inference of mechanisms and interactions. Additionally, we may need to refine expectations from the literature by considering that the effects of biotic and abiotic environmental factors occur in a multidimensional space in nature and may be interdependent [58]. It would be important to understand how many of our expectations are confined only to certain areas of this multidimensional space by, for example, leveraging on monitoring data to infer responses of organisms in natural conditions (as in this study) or designing factorial experiments that explore complex interactions among environmental variables [62]. Size is a master trait explaining emergent and aggregated ecosystems’ properties at equilibrium [5, 24]. We should ask to what extent can size provide understanding and prediction of short-term dynamics when systems are out-of-equilibrium [37, 63].

## Acknowledgements

This research was funded by the Swiss National Science Foundation (projects 182124 Aquascope and 202290 Cyanobloom) and the German Research Foundation (project 412375259 Aquascope), with contribution for infrastructure by the Swiss Federal Office for the Environment (contract Nr Q392-1149). F.P and P. I. also acknowledge the Eawag project #5221.00492.012.04. F.P., S.K. and M.B.-J. acknowledge the Eawag DF project Big-Data Workflow (#5221.00492.999.01). M.B.-J. Thanks to N. Schuwirth, E. Chollet-Ramampiandra, M. Hoege and A. Scheidegger for valuable conversations.

## Data Availability Statement

The data supporting this study’s findings, and the R scripts that generated the results, are openly available from the Eawag Research Data Institutional Repository (ERIC/Open) at https://doi.org/10.25678/000BV9. Raw abiotic data can be found at https://doi.org/10.25678/000C8P and raw biotic data can be found at https://doi.org/10.25678/000C2G.

## References

1. Falkowski PG, Katz ME, Knoll AH, Quigg A, Raven JA, Schofield O, et al. The evolution of modern eukaryotic phytoplankton. Science 7 2004; 305: 354–360.

2. Anneville O, Chang C-W, Dur G, Souissi S, Rimet F, Hsieh C-H. The paradox of re-oligotrophication: the role of bottom–up versus top–down controls on the phytoplankton community. Oikos 2019; 128: 1666–1677.

3. Rogers TL, Munch SB, Stewart SD, Palkovacs EP, Giron-Nava A, Matsuzaki S-IS, et al. Trophic control changes with season and nutrient loading in lakes. Ecol Lett 2020; 23: 1287–1297.

4. Merz E, Saberski E, Gilarranz LJ, Isles P, Sugihara G, Berger C, et al. Disruption of ecological networks in lakes by climate change and nutrient fluctuations. Nat Clim Chang 2023.

5. Wickman J, Litchman E, Klausmeier CA. Eco-evolutionary emergence of macroecological scaling in plankton communities. Science 2024; 383: 777–782.

6. Litchman E, Klausmeier CA. Trait-Based Community Ecology of Phytoplankton. Annual Review of Ecology, Evolution, and Systematics. 2008., 39: 615–639

7. Levine JM. Indirect facilitation: Evidence and predictions from a riparian community. Ecology 1999; 80: 1762–1769.

8. Shurin JB, Gruner DS, Hillebrand H. All wet or dried up? Real differences between aquatic and terrestrial food webs. Proceedings of the Royal Society B: Biological Sciences 2006; 273: 1–9.

9. Lima-Mendez G, Faust K, Henry N, Decelle J, Colin S, Carcillo F, et al. Determinants of community structure in the global plankton interactome. Science 2015; 348: 1262073.

10. Behrenfeld MJ, O’Malley R, Boss E, Karp-Boss L, Mundt C. Phytoplankton biodiversity and the inverted paradox. ISME Commun 2021; 1: 52.

11. Ehrlich E, Thygesen UH, Kiørboe T. Evolution of toxins as a public good in phytoplankton. Proceedings of the Royal Society B: Biological Sciences 6 2022; 289: 20220393.

12. Behrenfeld MJ, Boss ES. Student’s tutorial on bloom hypotheses in the context of phytoplankton annual cycles. Glob Chang Biol 2018; 24: 55–77.

13. Isles PDF, Pomati F. An operational framework for defining and forecasting phytoplankton blooms. Front Ecol Environ 2021; 19: 443–450.

14. Arteaga LA, Boss E, Behrenfeld MJ, Westberry TK, Sarmiento JL. Seasonal modulation of phytoplankton biomass in the Southern Ocean. Nat Commun 2020; 11: 5364.

15. Marañón E. Cell Size as a Key Determinant of Phytoplankton Metabolism and Community Structure. Annual Review of Marine Science. 2015., 7: 241–264

16. Andersen KH, Berge T, Gonçalves RJ, Hartvig M, Heuschele J, Hylander S, et al. Characteristic Sizes of Life in the Oceans, from Bacteria to Whales. Ann Rev Mar Sci 2016; 8: 217–241.

17. Gjoni V, Glazier DS, Wesner JS, Ibelings BW, Thomas MK. Temperature, resources and predation interact to shape phytoplankton size–abundance relationships at a continental scale. Glob Ecol Biogeogr 2023; 32: 2006–2016.

18. Pomati F, Shurin JB, Andersen KH, Tellenbach C, Barton AD. Interacting Temperature, Nutrients and Zooplankton Grazing Control Phytoplankton Size-Abundance Relationships in Eight Swiss Lakes. Front Microbiol 2020; 10: 3155.

19. Thomas MK, Fontana S, Reyes M, Kehoe M, Pomati F. The predictability of a lake phytoplankton community, over time-scales of hours to years. Ecology Letters. 2018., 21: 619–628

20. Orenstein EC, Ratelle D, Briseño-Avena C, Carter ML, Franks PJS, Jaffe JS, et al. The Scripps Plankton Camera system: A framework and platform for in situ microscopy. Limnol Oceanogr Methods 2020; 18: 681–695.

21. Merz E, Kozakiewicz T, Reyes M, Ebi C, Isles P, Baity-Jesi M, et al. Underwater dual-magnification imaging for automated lake plankton monitoring. Water Res 2021; 203: 117524.

22. Kyathanahally SP. Plankiformer. Github.

23. Kyathanahally SP, Hardeman T, Merz E, Bulas T, Reyes M, Isles P, et al. Deep Learning Classification of Lake Zooplankton. Front Microbiol 2021; 12: 746297.

24. Sprules WG, Barth LE. Surfing the biomass size spectrum: some remarks on history, theory, and application. Can J Fish Aquat Sci 2016; 73: 477–495.

25. Winslow L, Read J, Woolway R, Brentrup J, Leach T, Zwart J, et al. rLakeAnalyzer: Lake physics tools. R package version 1.11. 4.1. 2019.

26. Pomati F, Jokela J, Simona M, Veronesi M, Ibelings BW. An automated platform for phytoplankton ecology and aquatic ecosystem monitoring. Environ Sci Technol 2011; 45: 9658–9665.

27. Stockwell JD, Doubek JP, Adrian R, Anneville O, Carey CC, Carvalho L, et al. Storm impacts on phytoplankton community dynamics in lakes. Glob Chang Biol 2020; 26: 2756–2784.

28. Breiman L. Random Forests. Mach Learn 2001; 45: 5–32.

29. Bzdok D, Altman N, Krzywinski M. Points of Significance: Statistics versus machine learning. Nat Methods 2018; 15: 233–234.

30. Liaw A, Wiener M, Others. Classification and regression by randomForest. R news 2002; 2: 18–22.

31. Møller A, Jennions MD. How much variance can be explained by ecologists and evolutionary biologists? Oecologia 2002; 132: 492–500.

32. Greenwell B. Pdp: An R package for constructing partial dependence plots. R J 2017; 9: 421.

33. Rogers TL, Johnson BJ, Munch SB. Chaos is not rare in natural ecosystems. Nat Ecol Evol 2022; 6: 1105–1111.

34. Sommer U, Adrian R, Domis LDS, Elser JJ, Gaedke U, Ibelings BW, et al. Beyond the Plankton Ecology Group (PEG) Model: Mechanisms Driving Plankton Succession. Annu Rev Ecol Evol Syst 12 2012; 43: 429–448.

35. Benincà E, Huisman J, Heerkloss R, Jöhnk KD, Branco P, van Nes EH, et al. Chaos in a long-term experiment with a plankton community. Nature 2 2008; 451: 822–825.

36. Dakos V, Benincà E, van Nes EH, Philippart CJM, Scheffer M, Huisman J. Interannual variability in species composition explained as seasonally entrained chaos. Proc Biol Sci 8 2009; 276: 2871–2880.

37. Munch SB, Rogers TL, Johnson BJ, Bhat U, Tsai C-H. Rethinking the Prevalence and Relevance of Chaos in Ecology. Annu Rev Ecol Evol Syst 2022; 53: 227–249.

38. Ryabov AB, Rudolf L, Blasius B. Vertical distribution and composition of phytoplankton under the influence of an upper mixed layer. J Theor Biol 3 2010; 263: 120–133.

39. Fontana S, Jokela J, Pomati F. Opportunities and challenges in deriving phytoplankton diversity measures from individual trait-based data obtained by scanning flow-cytometry. Front Microbiol 2014; 5: 324.

40. Orenstein EC, Ayata S-D, Maps F, Becker ÉC, Benedetti F, Biard T, et al. Machine learning techniques to characterize functional traits of plankton from image data. Limnol Oceanogr 2022; 67: 1647–1669.

41. Collins S, Gardner A. Integrating physiological, ecological and evolutionary change: a Price equation approach. Ecol Lett 8 2009; 12: 744–757.

42. Enquist BJ, Norberg J, Bonser SP, Violle C, Webb CT, Henderson A, et al. Chapter Nine - Scaling from Traits to Ecosystems: Developing a General Trait Driver Theory via Integrating Trait-Based and Metabolic Scaling Theories. In: Pawar S, Woodward G, Dell AI (eds). Advances in Ecological Research. 2015. Academic Press, pp 249–318.

43. Winder M, Sommer U. Phytoplankton response to a changing climate. Hydrobiologia 2012; 698: 5–16.

44. Cloern JE. Why large cells dominate estuarine phytoplankton. Limnol Oceanogr 2018; 392–409.

45. Huisman J, Codd GA, Paerl HW, Ibelings BW, Verspagen JMH, Visser PM. Cyanobacterial blooms. Nat Rev Microbiol 2018; 16: 471–483.

46. Acevedo-Trejos E, Marañón E, Merico A. Phytoplankton size diversity and ecosystem function relationships across oceanic regions. Proceedings of the Royal Society B: Biological Sciences 2018; 285: 20180621.

47. Pierce RW, Turner JT. Ecology of Planktonic Ciliates in Marine Food Webs. Reviews in Aquatic Sciences 1992; 6: 139–181.

48. Verni F, Gualtieri P. Feeding behaviour in ciliated protists. Micron 1997; 28: 487–504.

49. Ger KA, Urrutia-Cordero P, Frost PC, Hansson LA, Sarnelle O, Wilson AE, et al. The interaction between cyanobacteria and zooplankton in a more eutrophic world. Harmful Algae 2016; 54: 128–144.

50. Gilbert JJ. Food niches of planktonic rotifers: Diversification and implications. Limnol Oceanogr 2022; 67: 2218–2251.

51. Wilke T, Ahlrichs WH, Bininda-Emonds ORP. The evolution of Synchaetidae (Rotifera: Monogononta) with a focus on Synchaeta : An integrative approach combining molecular and morphological data. J Zoolog Syst Evol Res 2020; 58: 823–857.

52. Stelzer C-P. Feeding behaviour of the rotifer Ascomorpha ovalis : functional response, handling time and exploitation of individual Ceratium cells. J Plankton Res 1998; 20: 1131–1144.

53. Stocker R, Seymour JR. Ecology and Physics of Bacterial Chemotaxis in the Ocean. Microbiol Mol Biol Rev 12 2012; 76: 792–812.

54. Sommer U, Stibor H, Katechakis A, Sommer F, Hansen T. Pelagic food web configurations at different levels of nutrient richness.pdf. Hydrobiologia 2002; 484: 11–20.

55. Zadereev E, Lopatina T, Ovchinnikov S, Tolomeev A. Grazing rate and feeding selectivity of small and large bodied cladocerans in waters from lakes with different salinity and phytoplankton structure. Journal of Oceanology and Limnology 2023.

56. Burkholder JM, Glibert PM. Eutrophication and Oligotrophication. In: Scheiner SM (ed). Encyclopedia of Biodiversity (Third Edition). 2024. Academic Press, Oxford, pp 442–478.

57. Brandl Z. Freshwater Copepods and Rotifers: Predators and their Prey. Hydrobiologia 2005; 546: 475–489.

58. Ye H, Sugihara G. Information leverage in interconnected ecosystems: Overcoming the curse of dimensionality. Science 2016; 353: 922–925.

59. Ryabov A, Kerimoglu O, Litchman E, Olenina I, Roselli L, Basset A, et al. Shape matters: the relationship between cell geometry and diversity in phytoplankton. Ecol Lett 2021; 24: 847–861.

60. Stanca E, Cellamare M, Basset A. Geometric shape as a trait to study phytoplankton distributions in aquatic ecosystems. Hydrobiologia 2013; 701: 99–116.

61. Padisák J, Soróczki-Pintér É, Rezner Z. Sinking properties of some phytoplankton shapes and the relation of form resistance to morphological diversity of plankton—an experimental study. Aquatic biodiversity 2003.

62. Thomas MK, Ranjan R. Designing More Informative Multiple-Driver Experiments. Ann Rev Mar Sci 2023.

63. Rogers TL, Munch SB, Matsuzaki S-IS, Symons CC. Intermittent instability is widespread in plankton communities. Ecol Lett 2023; 26: 470–481.

