## Supplementary information for "Automated plankton monitoring suggests a key role of microzooplankton and temperature for predicting dynamics of phytoplankton size classes"

#### Table of contents

**Supplementary Information** **1**
1. Supplementary Methods 2
Abiotic explanatory data: Nutrient data interpolation 2
2. Supplementary Results 3
3. Supplementary Tables 5
4. Supplementary Figures 10
5. References 24

### 1. Supplementary Methods

#### Abiotic explanatory data: Nutrient data interpolation

We used nitrate, ammonium and phosphate as measures of inorganic nutrient supply to the plankton community. Nutrient chemistry measurements were performed weekly, using standard colourimetric methods [1]. We predicted daily observations with a Random Forest model  (randomForest package V4.6.14 [2]). The models were trained on weekly nutrient measurements between 2019-2023. CTD and Meteo data from the same dates as chemistry measurements were used to train the weekly nutrient model. Where CTD or Meteo data were absent or contained more than one missing value, they were replaced with the values from the nearest available date within a ±3-day window, assuming similar environmental conditions.  Ammonium levels were best predicted with $O_2$ concentration, conductivity, air and water temperature, light penetration depth (depth at 5 PAR, **Fig. S2**) and Chlorophyll-a concentration. Nitrate levels used $O_2$ concentration, Chlorophyll-a concentration, water temperature, light penetration depth, pH and conductivity in the model. Phosphate was predicted using temperature, conductivity, pH, Chlorophyll-a concentration, $O_2$ concentration and light penetration depth. The models' performance on weekly observations was evaluated through 10-fold cross-validation, where 80% of the data were randomly sampled (without replacement) for training, and the remaining 20% were used for testing. The results of this evaluation are presented in **Table S4** and **Fig. S13**. Following validation and testing, the top-performing model for each nutrient was employed to predict its concentrations at a daily scale, utilising daily CTD profiles and meteorological measurements between the weekly data used for training (**Fig. S13**).

### 2. Supplementary Results

#### Water physics and chemistry

Abiotic environmental conditions followed a common seasonal pattern for a temperate lake, with high light irradiance of the photic zone from April to mid-August, while epilimnetic temperature ranged from a minimum of 7 °C in winter to a maximum of 23.8 °C in July. Thermocline depth ranged from 3.8 m in April to site depth (17.15 m) in winter due to deep mixing, and the depth of convective mixing ranged from 2.7 m in summer to 14.4 m in December. The stability of the water column increased from zero in April to a maximum in June and July. The depth of the thermocline and the mixed layer suggested high nutrient fluxes from the deep anoxic waters into the photic zone of the lake between August and mid-November. Ammonium and phosphate concentrations were low and stable until October when they increased due to autumn and winter mixing, while nitrate in contrast increased rapidly in April and decreased again throughout the year (**Fig. S2**).

#### Taxonomic composition of size bins

Bins one, two and three had similar taxonomic composition which visually differed from bins four, five and six. Bins five and six contained a more similar composition to each other than bin four across the entire time series and across time. The bin with the smallest size (size-bin one) was dominated by cryptophytes and smaller chlorophytes. This dominance was little variable across the study period. Its taxonomic composition was very similar to the composition of the second and third smallest sized bins. In comparison to bin one, the second and third smallest bins contained an increasing proportion of *Chroococcales*, chrysophytes (*Dinobryon*, *Mallomonas*), dinoflagellates, diatoms (centric and pennates) and colonial cyanobacteria. The proportion of colonial cyanobacteria in bins one to three was much lower than in the larger size-bins (four to six). Compared to bins one and two, the third smallest bin had a temporally more variable taxonomic composition. Bins four to six contained an increasing proportion of large diatoms (*Asterionella*, *Aulacoseira*, *Fragilaria*) and the dinoflagellate *Ceratium sp*. with bin four containing the largest proportion of centric diatoms and intermediate-sized dinoflagellates across all bins. Compared to the two largest bins, bin four contained a larger proportion of large chlorophytes. Bins four and five contained the largest proportions of colonial cyanobacteria. The largest bin further contained large cells (*Synedra acus var. angustissima*) and colonies such as *Aulacoseira sp.*, *Dinobryon sp.* and *Uroglena sp*. (**Fig. 1, Fig. S1, Fig. S7, Tab. S3**).

We note that we use the area of identified objects to create the above size-bins, as opposed to many previous studies that use cell volume in $\mu m^3$ [3–6], which is generally nominal (not measured for each individual object, but assumed based on species average dimensions and shape). The use of area as opposed to volume to bin phytoplankton objects should not change the composition and dynamics of the size bins reported above.

With our instrument, we obtain the area of each single identified object (based on the binary mask created by the edge detector), and the maximum and minimum length of the object (https://github.com/tooploox/SPCConvert). By assuming that the third (unseen) dimension of each object is the same as the minimum observed dimension in a 2D plane, we could estimate an approximate biovolume, which would match the current standard in the field - this calculation is already implemented in the image processing pipeline SPCConvert. We however know that the assumption of the third dimension is heavy, and so far used the area of the object instead as an unbiased measure of body size [7, 8], since this allows us to account for the complex structure of the phytoplankton and avoid potential untested assumptions regarding the third unseen dimension.

Area scales with volume with an exponent of 2/3 (area scales by a factor of $k^2$ while volume scales by a factor of $k^3$ with changes in length, width, and height of an object). If an object doubles its length, width, and height, its area will quadruple ($2^2 = 4$), and its volume will increase by a factor of eight ($2^3 = 8$). If you halve the dimensions of an object, its area will become one-fourth ($0.5^2 = 0.25$), and its volume will become one-eighth ($0.5^3 = 0.125$). These variations will not change the definition and composition of our size bins, which were built on a Log10 scale and were equally spaced, nor their temporal changes in densities.

### 3. Supplementary Tables

**Table S1**: Size range of the six bins used in this analysis and RF model performance. Bins were generated on a Log10 scale of area. Mean major and minor axis length are also displayed, as well as the Pseudo-$R^2$ values that were extracted from the RF models and represent the variance explained.

| **Size-bin** | **Area range** | **Mean major axis length ± SD** | **Mean minor axis length ± SD** | **Model performance (Pseudo-$R^2$)** |
|---|---|---|---|---|
| 1 | < 239.88 $\mu m^2$ | 17.7 ± 3.54 µm | 12.56 ± 1.81 µm | 68.52 |
| 2 | (239.88 $\mu m^2$, 501.19 $\mu m^2$] | 28.1 ± 7.85 µm | 17.31 ± 3.18 µm | 63.46 |
| 3 | (501.19 $\mu m^2$, 1047.13 $\mu m^2$] | 40.96 ± 12.91 µm | 24.07 ± 5.44 µm | 62.28 |
| 4 | (1047.13 $\mu m^2$, 2187.76 $\mu m^2$] | 65.84 ± 27.58 µm | 34.38 ± 8.97 µm | 66.6 |
| 5 | (2187.76 $\mu m^2$, 4570.88 $\mu m^2$] | 124.22 ± 58.39 µm | 46.37 ± 18.08 µm | 61.36 |
| 6 | > 4570.88 $\mu m^2$ | 345.51 ± 251.08 µm | 65.4 ± 40.75 µm | 61.43 |

**Table S2**: Zooplankton taxa generated by a CNN classifier [9] were summed into six classes. Displayed are all taxa from the CNN classifier used for this grouping and the magnification at which the taxa were imaged.

| Zooplankton class | Taxa included from CNN classifier | Broad grazer group | magnification |
|---|---|---|---|
| Daphnids | - Bosmina<br>- Daphnia<br>- Diaphanosoma | Mesozooplankton | 0p5x |
| Cyclopoid copepods | - Cyclops | Mesozooplankton | 0p5x |
| Calanoid copepods | - Eudiaptomus | Mesozooplankton | 0p5x |
| Nauplii | - Nauplius | Microzooplankton | 0p5x |
| Rotifers | - Asplanchna<br>- Conochilus<br>- Kellicottia<br>- Keratella cochlearis<br>- Keratella quadrata<br>- Polyarthra<br>- Rotifers<br>- Synchaeta<br>- Trichocerca | Microzooplankton | 0p5x |
| Ciliates | - Askenasia<br>- Ciliate_round<br>- Ciliates<br>- Ciliates_blue<br>- Ciliates_green<br>- Coleps<br>- Didinium<br>- Paradileptus<br>- Strombidium<br>- Tintinidium<br>- Tintinopsis<br>- Vorticella_epistylis_like | Microzooplankton | 5p0x |

**Table S3**: Phytoplankton taxa generated by a CNN classifier [9] were summed into broader taxonomic classes. Displayed are all taxa from the CNN classifier used for this grouping.

| **Phytoplankton class** | **Taxa included from CNN classifier** |
|---|---|
| Asterionella | - Asterionella_colonies<br>- Asterionella_partial |
| Aulacoseira | - Aulacoseira |
| Centric Diatoms | Centric_diatom |
| Ceratium | Ceratium |
| Chlorophyte | - Coelastrum_reticulatum<br>- Elakatothrix<br>- Planktosphaeria |
| Chlorophyte Big | - Chlorophyte_colonial_dividing<br>- Chlorophyte_square<br>- Gonium<br>- Hormidium_like<br>- Pandorina<br>- Scenedesmus |
| Chlorophyte Small | - Chlorophyte<br>- Chlorophyte_elongated<br>- Chlorophyte_frame<br>- Oocystaceae<br>- Phacotus<br>- Tetraedron |
| Chroococcales | Chroococcales |
| Closterium | Closterium |
| Cosmarium | Cosmarium |
| Cryptophyte | - Cryptomonas_cryptophyceae<br>- Cryptophytes_blurry<br>- Rhodomonas |
| Cyanobacteria_colonial | - Coelosphaerium<br>- Cyanobacteria_colonial_blue<br>- Cyanobacteria_colonial_clathrate<br>- Cyanobacteria_colonial_probably<br>- Gomphosphaeria |
| Cyanobacteria_filament | - Cyanobacteria_filamentous<br>- Dolichospermum<br>- Limnoraphis<br>- Planktothrix |
| Diatom | Plankton_z |

| | |
|---|---|
| Dinobryon | - Dinobryon<br>- Dinobryon_single_cell |
| Dinoflagellate | - Dinoflagellate_diamond<br>- Gymnodinium |
| Filament | Filament |
| Fragilaria | Fragilaria |
| Mallomonas | - Mallomonas_akrokomos<br>- Mallomonas_big |
| Pediastrum | Pediastrum |
| Pennate Diatom | - Pennate_diatom<br>- Synedra |
| Peridinium | Peridinium |
| Staurastrum | Staurastrum |
| Synedra angustissima | Synedra_angustissima |
| Synura | Synura |
| Uroglena | Uroglena |

**Table S4:** Random Forest Model Performance for weekly prediction of nutrient concentration in lake Greifensee for the period 2019-2023.

| | Cross Validation | | | | Testing | |
|---|---|---|---|---|---|---|
| | Mean RMSE[2] | SD[1] RMSE[2] | Mean R-squared | SD[1] R-squared | RMSE[2] | R-squared |
| Ammonium µg/L | 37.974 | 13.531 | 0.678 | 0.248 | 43.536 | 0.551 |
| Nitrate µg/L | 0.211 | 0.047 | 0.670 | 0.143 | 0.166 | 0.745 |
| Phosphate µg/L | 4.801 | 1.616 | 0.854 | 0.120 | 6.328 | 0.600 |

1. SD: Standard Deviation; 2. RMSE: Root Mean Squared Error

### 4. Supplementary Figures

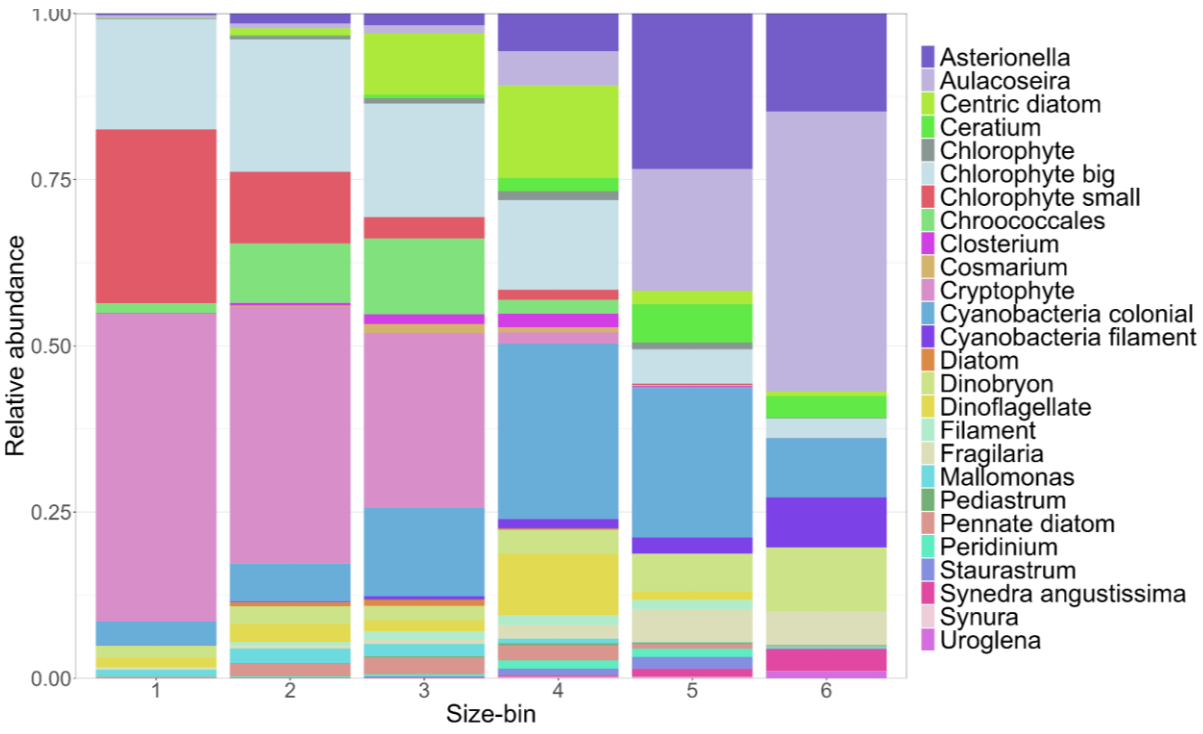


**Fig. S1.** Relative abundance of taxonomic classes per size bin. Classes are aggregated into broader taxonomic levels for easier display. Some classes contain more taxonomic groups than others. E.g. *Synedra angustissima* has 1 species while cryptophytes include all cryptomonas sp., rhodomonas sp. and other cryptophytes.

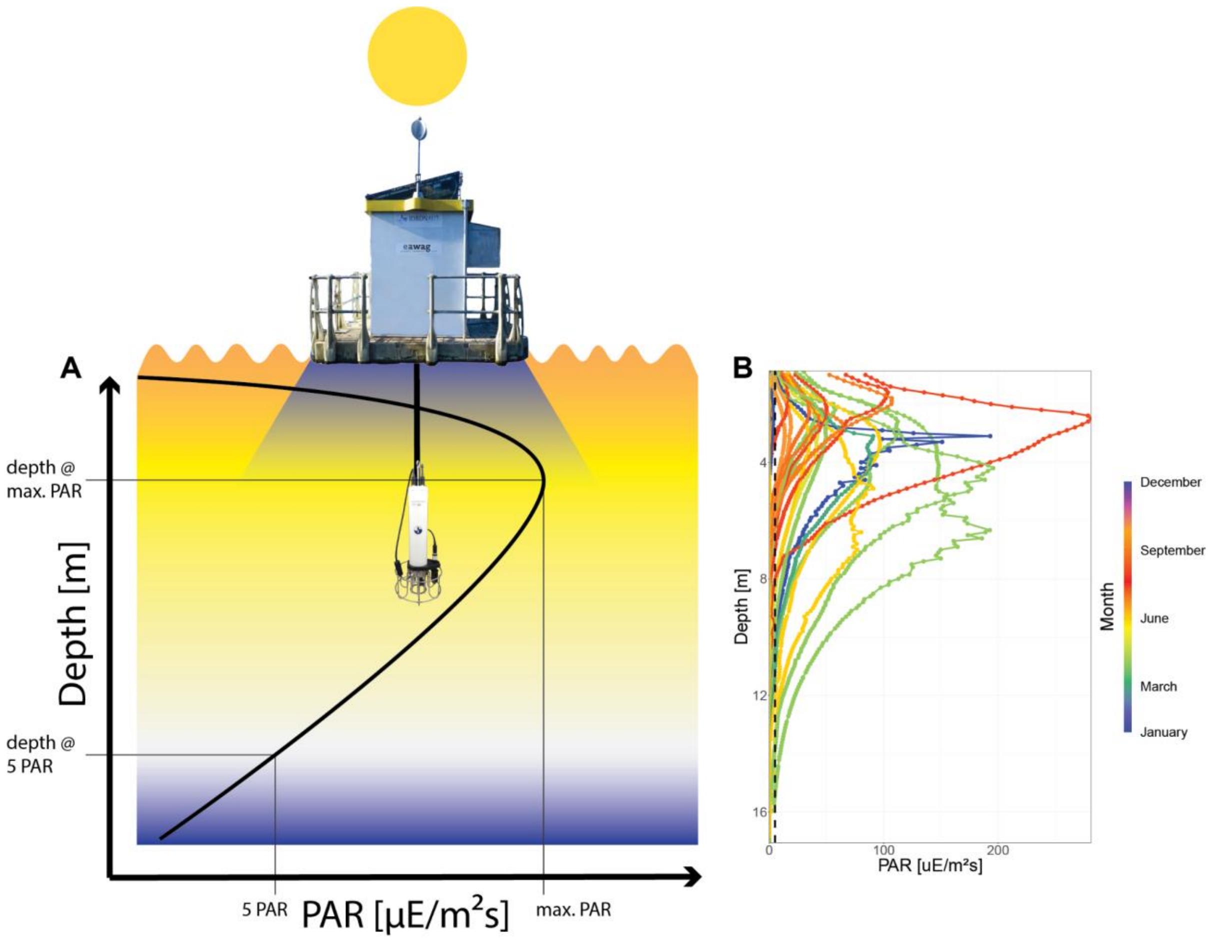


**Fig. S2**: A) Depth at 5 PAR (light penetration depth) is extracted from PAR profiles excluding all depths, where the platform shades the sensor. B) A subset of PAR profiles across the 4 years of data with a vertical dashed line at 5 PAR. The depth at max PAR varies across the season, years and across time of the day.

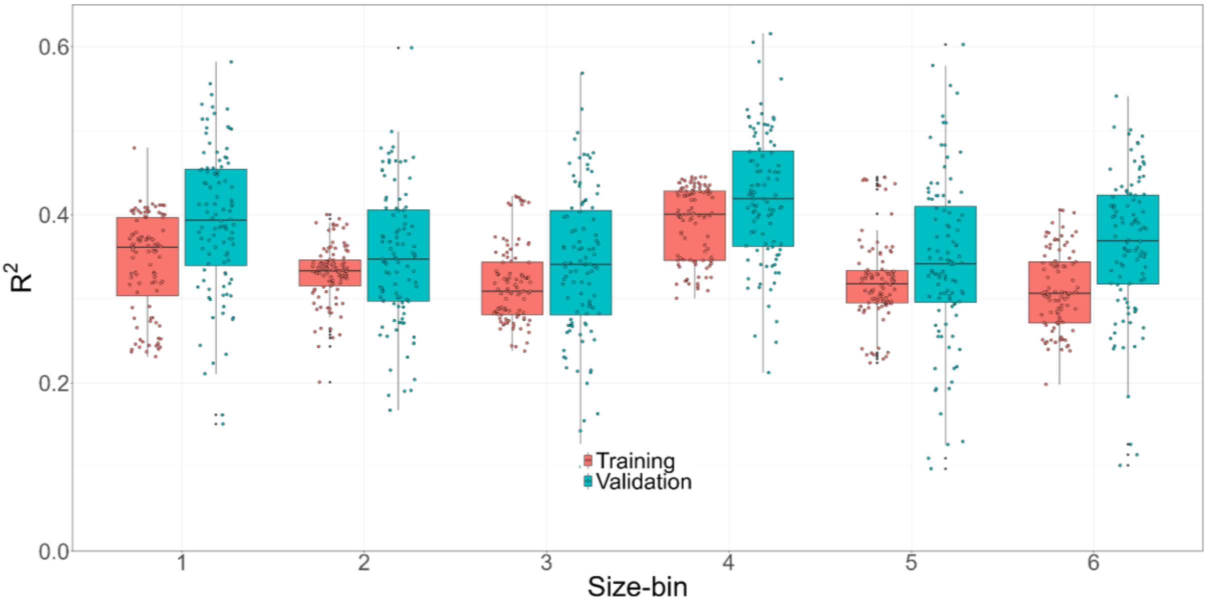


**Fig. S3**: $R^2$ from a linear model fit between the observed and the predicted accumulation rate from a RF model based on 90% of the data in a random split ten-fold cross validation that was repeated 10 times. Training $R^2$ in red was extracted from the RF model and validation $R^2$ in blue was extracted from linear models as described above.

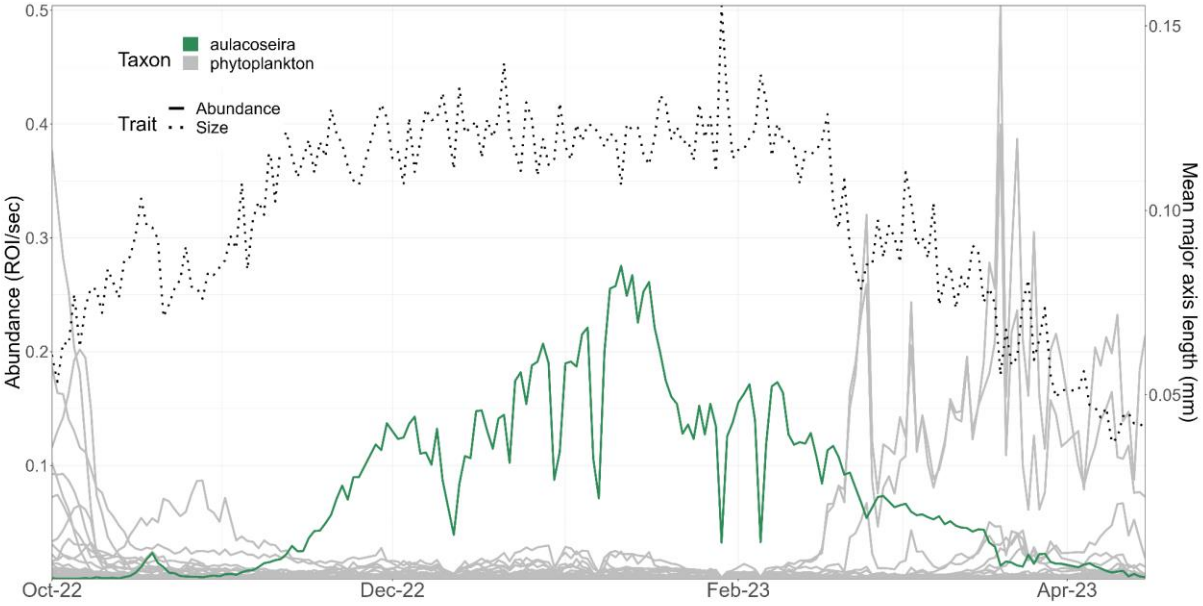


**Fig. S4**: *Aulacoseira sp.* bloom in winter 2022/2023. Abundance of Aulacoseira (green) compared to the rest of the phytoplankton taxa (gray) and the overall community size (mean major axis length, dotted line).

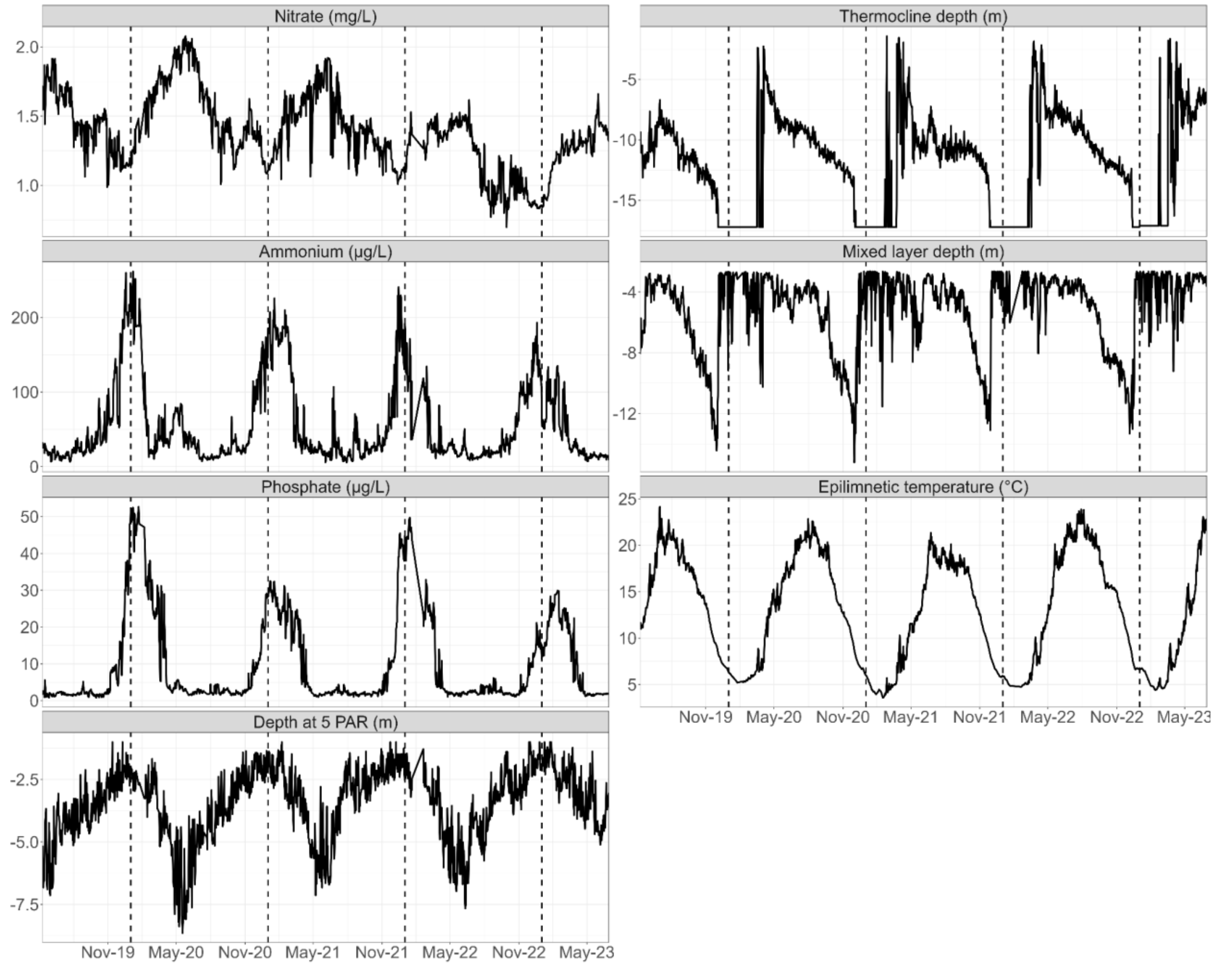


**Fig. S5.** Time series of abiotic explanatory variables. Nutrients (ammonium, nitrate, phosphate) were predicted from weekly measurements to daily observations (see SI Methods). Mixed layer depth, thermocline depth and epilimnetic temperature were calculated with the R package rLakeAnalyzer. Depth at 5 PAR was extracted from automated CTD profiles (Fig. S2). Note that in winter, Greifensee is mixed and there is no thermocline. Therefore, thermocline depth was set to the site depth (17m). In spring, thermocline depths are unstable until they stabilise.

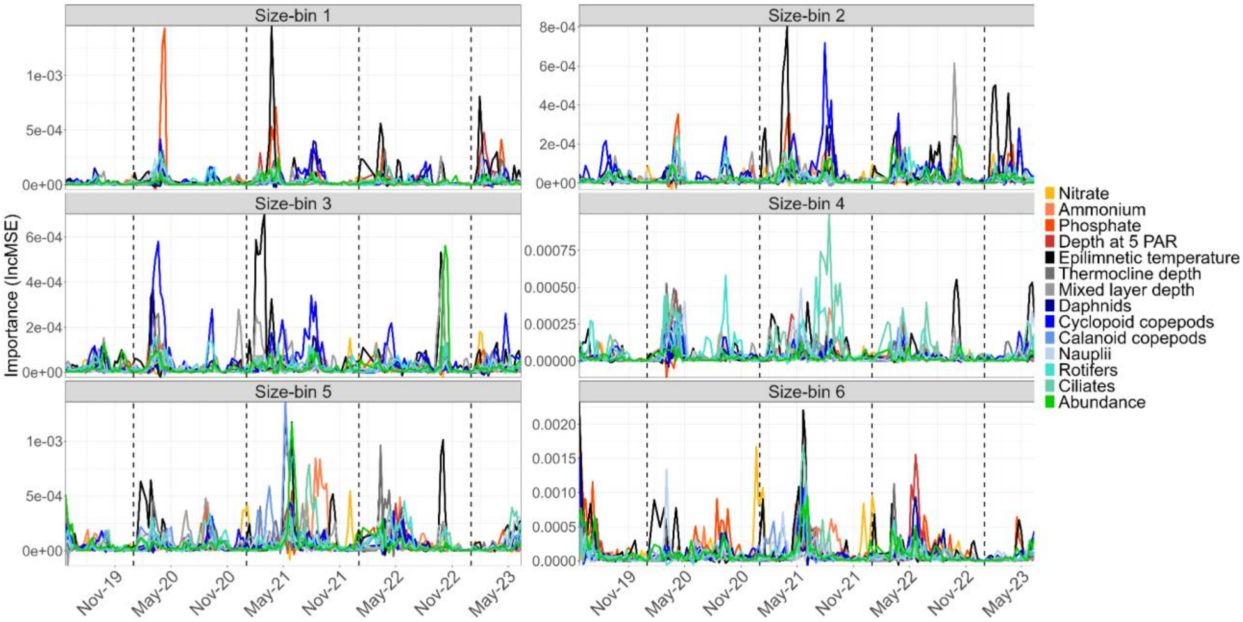


**Fig. S6.** Importance (IncMSE) of all variables across time. Mid-dates from a 28-day moving window are reported. Each panel represents one bin.

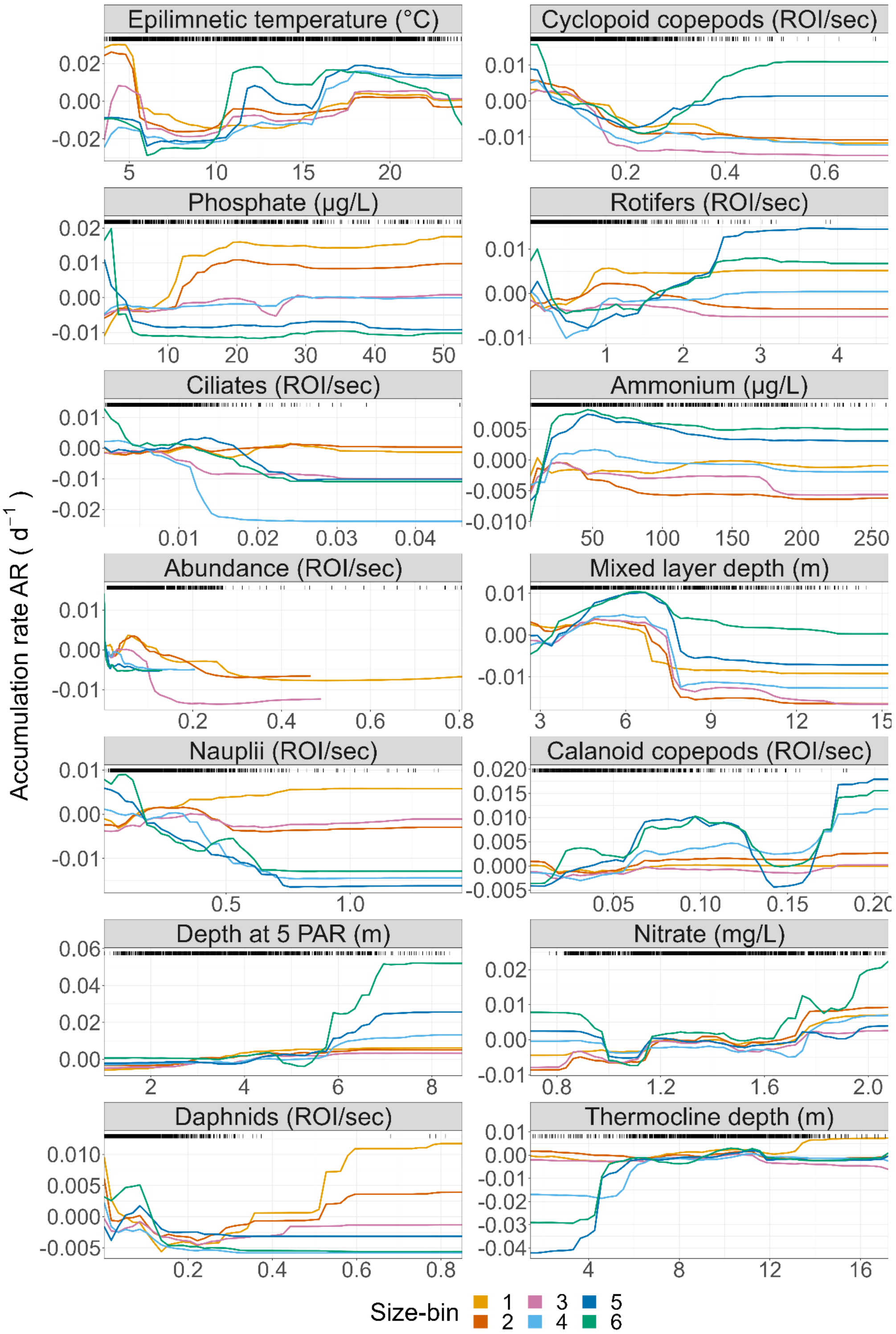


**Fig. S7:** Partial-effects of all predictors across all size-bins: these dependencies emerge from predicting the response variable (AR) over varying levels of the chosen explanatory variable (X-axis, in the plot title) while holding the other variables in the RF model at their medians. Each line (colour) represents one size-bin. The distribution of the model-training data is shown as black bars on top of each plot.

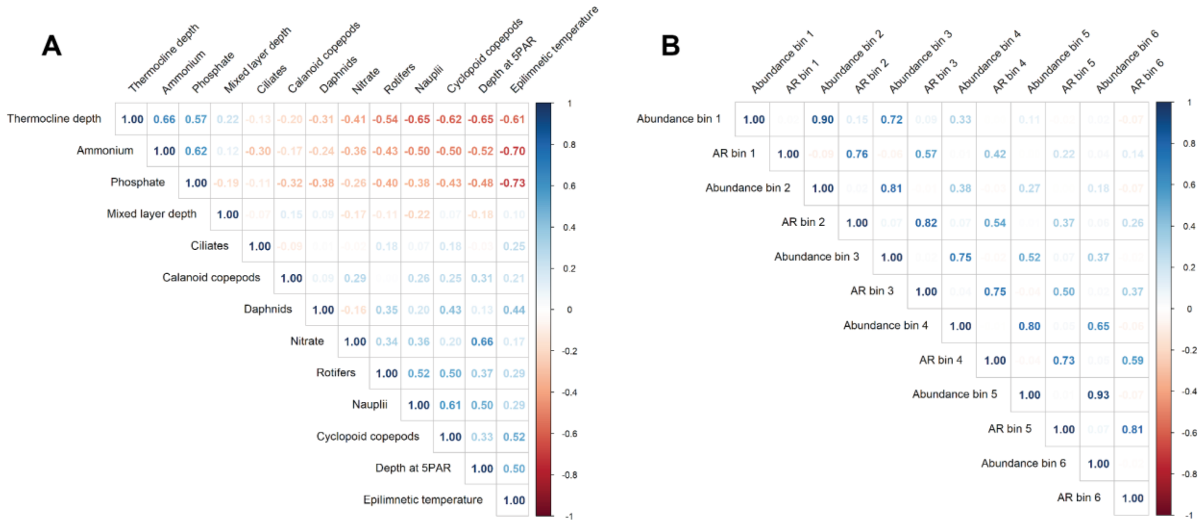


**Fig. S8:** Spearman correlation of **A)** explanatory biotic and abiotic variables and **B)** abundance and AR of each size-bin in the field data analysis

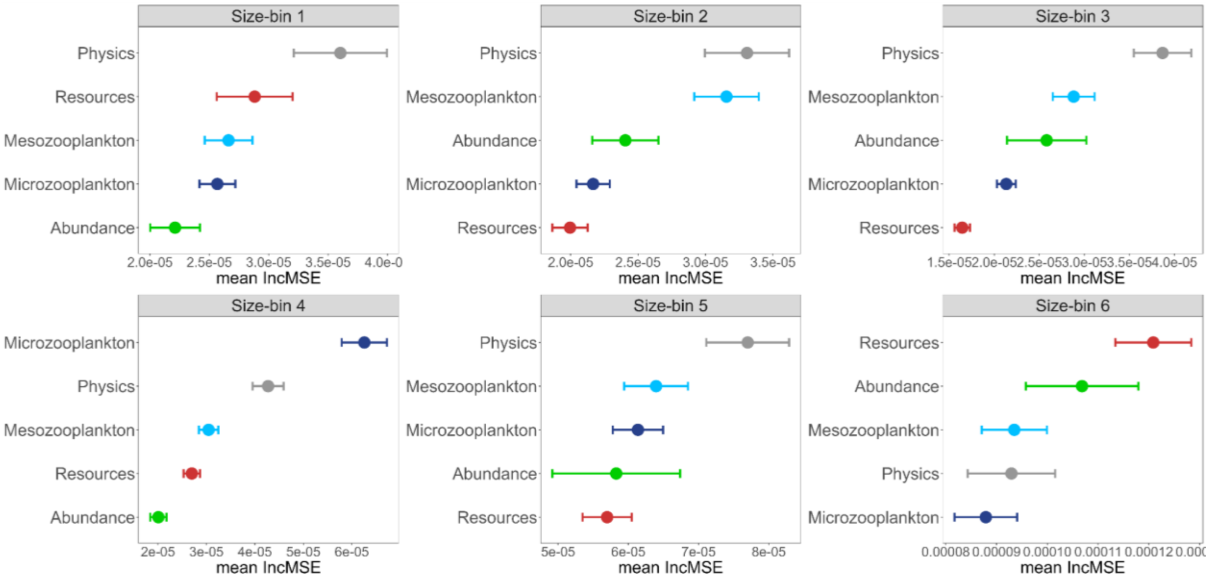


**Fig. S9:** Overall mean importance (IncMSE) ± standard error of each explanatory variable group across the entire time series (based on 28 d moving windows) for each size bin. In color, we display the predictor group the variable belongs to (colour coding as in Fig. 3, main text).

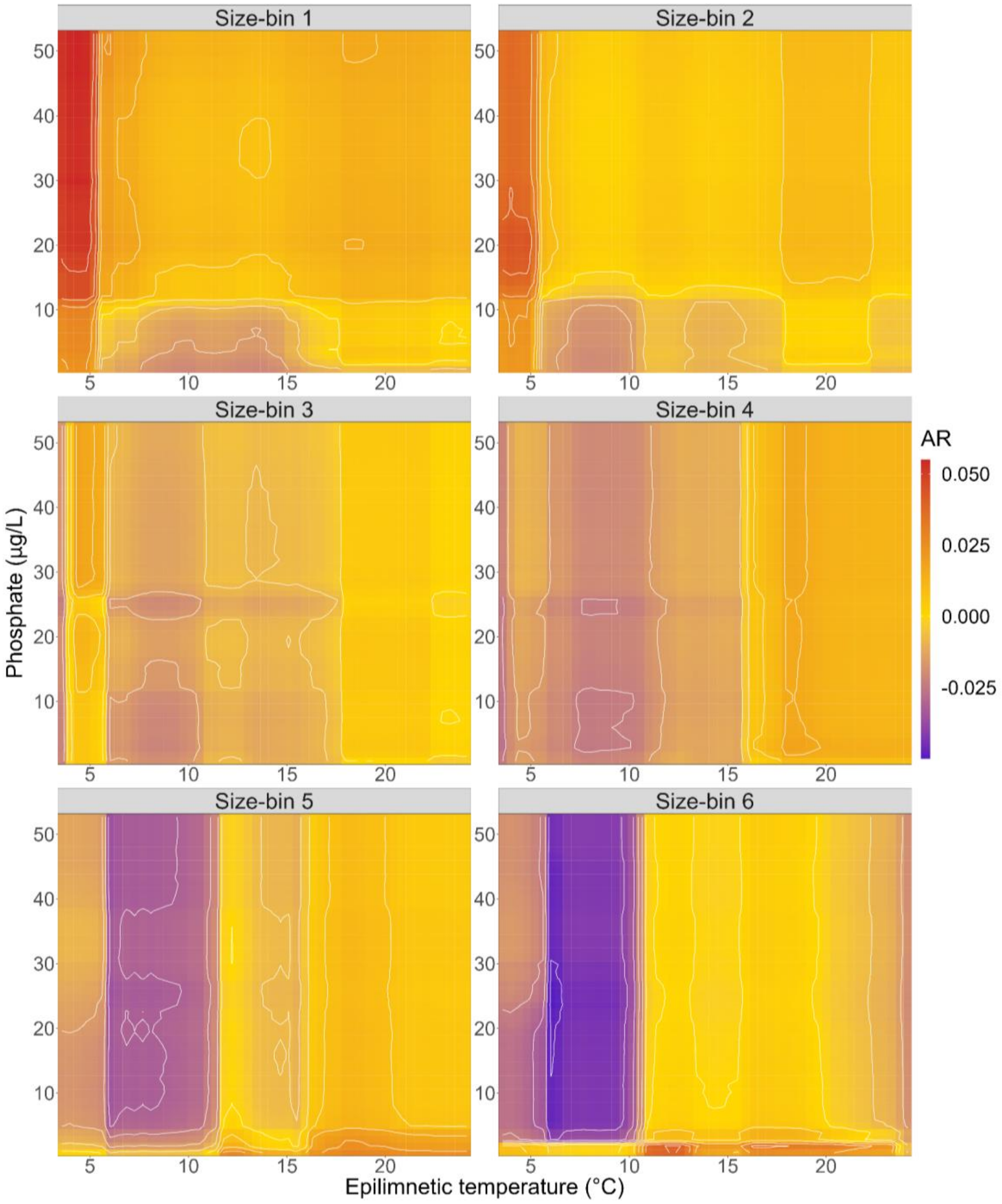


**Fig. S10:** Partial effects of epilimnetic temperature (x-axis) and phosphate (y-axis) across all size-bins on AR (in colour): these dependencies emerge from predicting the response variable (AR) over varying levels of the chosen explanatory variables while holding the other variables in the RF model at their medians.

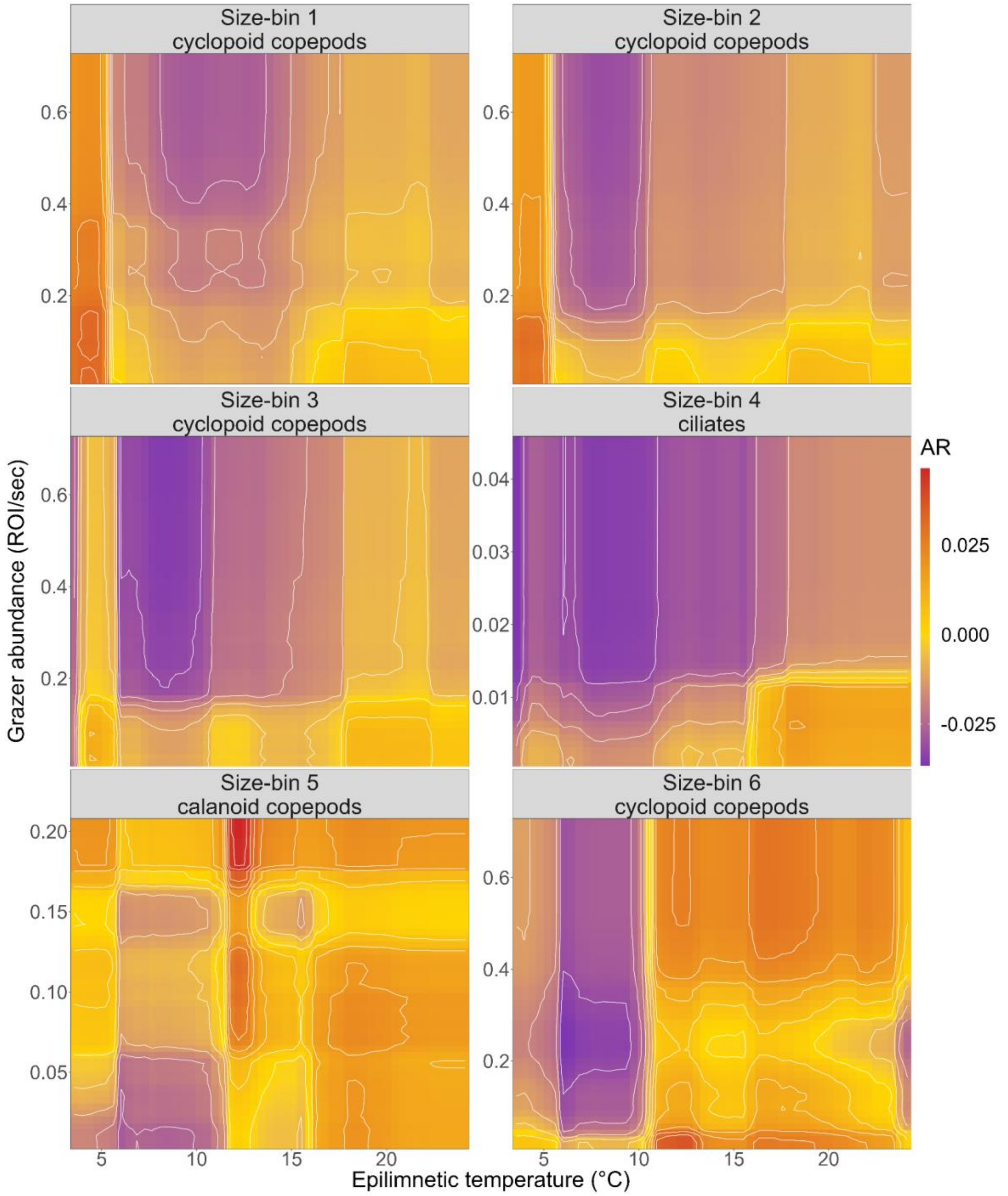


**Fig. S11**: Partial effects of epilimnetic temperature (x-axis) and most important grazer abundance (y-axis) across all size-bins on AR (in colour): these dependencies emerge from predicting the response variable (AR) over varying levels of the chosen explanatory variables while holding the other variables in the RF model at their medians. The grazer was chosen from Fig. 3 as the grazer variable with the highest overall mean importance (IncMSE) over the entire study period for each size-bin.

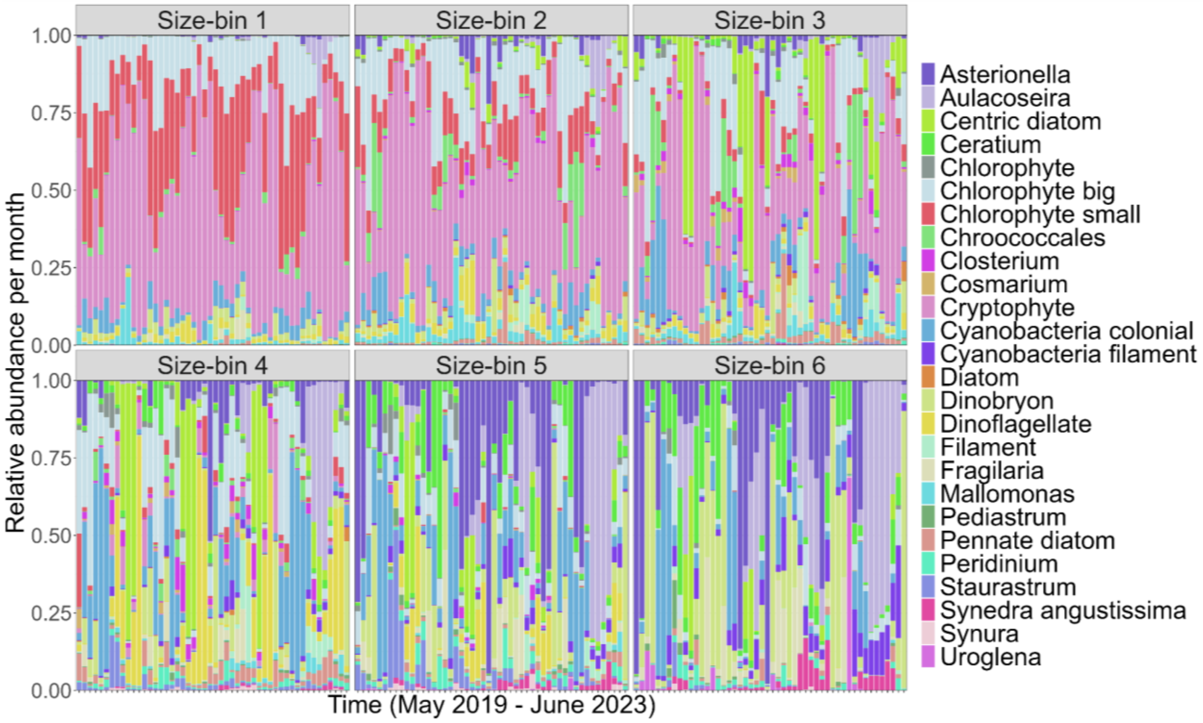


**Fig. S12:** Changing taxonomic composition and relative abundance per taxonomic group across time and size-bin. Abundances were summed across each month of the study period. The taxonomic composition of the smaller size-bins is less variable across time compared to the larger size-bins.

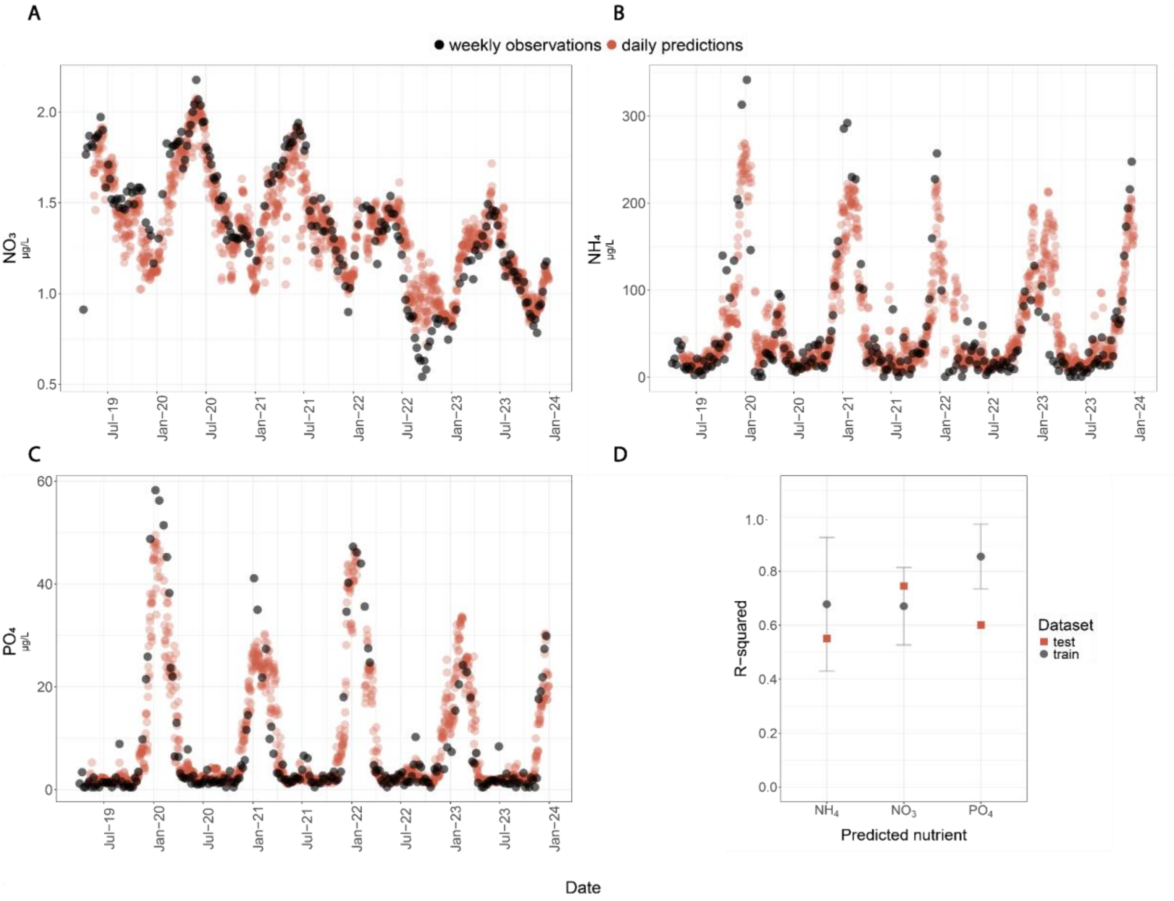


**Fig. S13**: Weekly and Daily Nutrient Predictions: Model Evaluation. (A, B, C) Daily predictions (red) from models trained on weekly observations compared to actual weekly observations (grey). The predictions consistently fall within the range of the weekly observations for all models. (D) Model performance measured by R-squared during training (grey) and testing (red). Error bars represent the standard deviation in training, calculated using 10-fold cross-validation. Testing was conducted once.

### 5. References


1. Awwa, Wef. Standard Methods for the examination of water and wastewater. 1999.
2. Liaw A, Wiener M, Others. Classification and regression by randomForest. *R news* 2002; **2**: 18–22.
3. Marañón E. Cell Size as a Key Determinant of Phytoplankton Metabolism and Community Structure. *Ann Rev Mar Sci* 2015; **7**: 241–264.
4. Hillebrand H, Acevedo-Trejos E, Moorthi SD, Ryabov A, Striebel M, Thomas P, et al. Cell size as driver and sentinel of phytoplankton community structure and functioning. *Funct Ecol* 2021.
5. Ryabov A, Kerimoglu O, Litchman E, Olenina I, Roselli L, Basset A, et al. Shape matters: the relationship between cell geometry and diversity in phytoplankton. *Ecol Lett* 2021; **24**: 847–861.
6. Wickman J, Litchman E, Klausmeier CA. Eco-evolutionary emergence of macroecological scaling in plankton communities. *Science* 2024; **383**: 777–782.
7. Merz E, Kozakiewicz T, Reyes M, Ebi C, Isles P, Baity-Jesi M, et al. Underwater dual-magnification imaging for automated lake plankton monitoring. *Water Res* 2021; **203**: 117524.
8. Merkli S, Reyes M, Pomati F. Laboratory application of the Aquascope approach of automated imaging and classification for long-term plankton monitoring. *bioRxiv* . 2024. , 2024.02.23.581739
9. Kyathanahally SP. Plankiformer. Github. https://github.com/kspruthviraj/Plankiformer
